# FOXP2-immunoreactive, corticothalamic pyramidal cells in neocortical layers 6a and 6b are tightly regulated by neuromodulatory systems

**DOI:** 10.1101/2024.06.21.599838

**Authors:** Guanxiao Qi, Danqing Yang, Fernando Messore, Arco Bast, Felipe Yáñez, Marcel Oberlaender, Dirk Feldmeyer

## Abstract

The *FOXP2*/*Foxp2* gene is involved in fine motor control in many vertebrate species; in humans, it is one of the candidate genes thought to play a prominent role in language production. Several studies suggest that in the neocortex, *Foxp2* is exclusively expressed in a subset of corticothalamic (CT) pyramidal cells (PCs) in layer 6 (L6). However, the morphological and intrinsic electrophysiological, synaptic and neuromodulatory properties of *Foxp2*-expressing L6 PCs remain largely unknown. Here we systematically characterise these properties for FOXP2-positive (FOXP2+) PCs across L6 in the rat somatosensory cortex. We find that L6 FOXP2+ PCs are distinct in all of these properties from those of L6 FOXP2-negative (FOXP2–) neuronal cell types. We show that L6 FOXP2+ PCs project exclusively to thalamus. In upper L6 (L6a), FOXP2+ PCs innervate either the first-order thalamus or both first and higher-order thalamic nuclei. FOXP2+ PCs in deep L6 (L6b) project almost exclusively to higher-order thalamus. Synaptic connections established by L6a and L6b FOXP2+ PCs exhibit low synaptic release probability, whereas L6 corticocortical PCs have a high release probability. Both L6a and L6b FOXP2+ PCs respond strongly to acetylcholine (ACh), which in the absence of TTX results in firing of action potential (AP) trains. Notably, L6b but not L6a FOXP2– PCs are highly sensitive to ACh. In addition, L6b FOXP2+ PCs close to the white matter border show strong responses to dopamine that develop into prolonged AP firing. Our data suggest that FOXP2 is a marker for CT PCs in L6 that are strongly controlled by neurotransmitters such as ACh and dopamine. These findings are in line with a pivotal role for both L6a and L6b CT PCs as modulators of thalamic activity.

## Introduction

The forehead box transcription factor P2 (FOXP2) gene (in humans FOXP2, in rodents Foxp2) is a highly conserved transcription factor present across many vertebrate species. It plays a crucial role in the neuronal circuits controlling vocalization and language, particularly in mammals and birds (Bolhuis et al., 2010; Lai et al., 2001; Vargha-Khadem et al., 2005). Mutations in the FOXP2 gene are associated with defective speech development and significant difficulties in controlling vocal-motor actions (for reviews see Enard, 2011; Graham and Fisher, 2015; Lai et al., 2001; Vargha-Khadem et al., 2005; Vernes et al., 2011; Watkins et al., 2002). Additionally, in humans, FOXP2 has been implicated in various neuropsychiatric disorders such as autism spectrum disorder, attention-deficit hyperactivity disorder (ADHD), and schizophrenia (e.g. Co et al., 2020; Demontis et al., 2019; Konopka and Roberts, 2016; Li and Pozzo-Miller, 2020; Meyer et al., 2022; Mukamel et al., 2011; Walker et al., 2012). These disorders may result from dysfunctional neuronal networks caused by disruptions in the interaction between FOXP2 and its transcription targets..

FOXP2 regulates the transcription of numerous downstream target genes (den Hoed et al., 2021; Konopka et al., 2012; Spiteri et al., 2007). It is involved in several neurodevelopmental processes, including neurite outgrowth, neuronal subtype specification, and the formation of synaptic circuits (Chen et al., 2016; Chiu et al., 2014; Reimers-Kipping et al., 2011; Sia et al., 2013; Tsui et al., 2013; Vernes et al., 2011). Moreover, *Foxp2* is critical for thalamocortical patterning and the proper formation of thalamocortical projections (Antón-Bolaños et al., 2019; Ebisu et al., 2017). While *Foxp2* deletion in the neocortex does not lead to obvious histoarchitectural deficits, it does result in behavioral impairments, such as difficulties in motor skill learning, which may indicate impaired neuronal microcircuit formation (French et al., 2019; Kast et al., 2019; Medvedev et al., 2018). In humans, single nucleotide polymorphisms in FOXP2 have been linked to ADHD, potentially due to synapse formation deficits (Demontis et al., 2019).

*Foxp2* is expressed in almost all neural networks involved in sensorimotor integration, modulation, and feedback control of fine motor output, including the olivo-cerebellar loop and cortico-basal ganglia networks, and is also found in neurons of several thalamic nuclei and all neocortical areas (Chen et al., 2016; Foster et al., 2021; Gao et al., 2018; Milardi et al., 2019). *Foxp2* expression is layer- and neuronal cell type-specific and restricted to a subset of excitatory projection neurons that are almost exclusively located layers 6a (L6a) and 6b of rodent neocortex (Ferland et al., 2003; Hisaoka et al., 2010; Ma et al., 2022). In layer 6a, *Foxp2* expression is indicative of corticothalamic (CT) projection neurons (Co et al., 2020; Druart et al., 2020; Kast et al., 2019; Sundberg et al., 2017) that co-express the neurotensin receptor 1 (*Ntsr1*) gene. Transcriptomic data suggest the presence of multiple L6 *Foxp2*-expressing glutamatergic neuron types (Tasic et al., 2018). In certain neocortical areas, particularly in the premotor and medial motor cortices, *Foxp2* is also found in a subset of deep L5 pyramidal cells (PCs), albeit at low density, which show, however, pyramidal tract projections (Campbell et al., 2009; Hisaoka et al., 2010; Kast et al., 2019). Furthermore, in mouse medial prefrontal cortex (mPFC), at least a subset of *Foxp2*-expressing L6 PCs projects to the ventral tegmental area, which in turn sends dopaminergic afferents to the neocortex and is involved in in numerous cognitive processes including decision making and working memory (Babiczky and Matyas, 2022).

The expression of *Foxp2* is tightly regulated by neuromodulators such as acetylcholine (ACh) and dopamine. *Foxp2* expression coincides with the expression of the *Chrna5* gene, which encodes the α_5_ subunit of the nicotinic acetylcholine receptors (nAChRs) and selectively enhances the activity of L6 CT neurons (Bailey et al., 2012; Kassam et al., 2008; Venkatesan and Lambe, 2020). The nAChR α_5_ subunit conveys increased ligand efficacy, increased Ca^2+^ permeability and reduced receptor desensitisation suggesting that L6 CT neurons exhibit a pronounced nAChR response (Kuryatov et al., 2008; Scholze and Huck, 2020). It has also been proposed that type 1 dopamine receptors regulate the activity of L6 CT PCs (Gaspar et al., 1995). Furthermore, studies have demonstrated reduced expression of the dopamine receptor 1 gene (*Drd1*) in neonate and adult *Foxp2* knockout mice in excitatory and inhibitory neurons in the frontal cortex (Co et al., 2020). This suggests that *Foxp2* may play a significant role in the development of dopamine-modulated cortical circuits. Hence, the CT microcircuitry in cortical L6 is regulated by at least two neuromodulatory systems involved in motor control(Speranza et al., 2021) and motor learning (Li and Hollis, 2021).

The present study aimed to investigate the morphology, electrophysiology, synaptic properties, and neuromodulation of both FOXP2-immunopositive (FOXP2+) and FOXP2-immunonegative (FOXP2-) PCs in the rat primary somatosensory (S1) cortex. In layer 6, FOXP2+ PCs exhibited three distinct CT projection patterns: a projection solely to the first-order ventral posterior medial thalamic nucleus (VPM), a projection to both VPM and the higher-order posterior medial complex (POm), and a projection solely to the POm. Almost all POm-projecting neurons were located in deep layer 6, i.e., layer 6b. Corticocortical (CC) excitatory neurons in layer 6a and 6b were consistently FOXP2–. Furthermore, cholinergic and dopaminergic neuromodulation of L6a and L6b excitatory neurons correlates with the Foxp2 expression pattern and the neuronal location within L6. Our findings indicate that FOXP2 is a marker for L6a and L6b CT PCs whose output is under tight control of cholinergic and dopaminergic neuromodulatory systems. These systems play crucial roles in motor behaviour, arousal, and vigilance.

## Results

### Expression of FOXP2 in cortical and subcortical areas

In the neocortex, the expression of the *Foxp2* gene is a highly specific marker of cortical layer 6 (Ferland et al., 2003; Hisaoka et al., 2010). Here, FOXP2 immunoreactivity was analysed across cortical and subcortical areas in 150 µm thick thalamocortical slices. FOXP2-related immunofluorescence was found exclusively and uniformly in layer 6 of the entire neocortex including the peri- and entorhinal cortex (Co et al., 2020; Druart et al., 2020; Kast et al., 2019) (**Figure 1**). In adjacent cortical areas, *i.e.* the secondary somatosensory cortex (S2) and primary motor cortex (M1) FOXP2 labelling was predominantly observed in layer 6 and to a markedly lesser extent in deep layer 5 (**Figure 1A,B**). In subcortical areas, the FOXP2 signal was robust in the striatum and in particular in the POm (**Suppl. Figure S1**), but only scattered in the VPM, and virtually absent in the hippocampus (**Figure 1A**).

**Figure 1.**
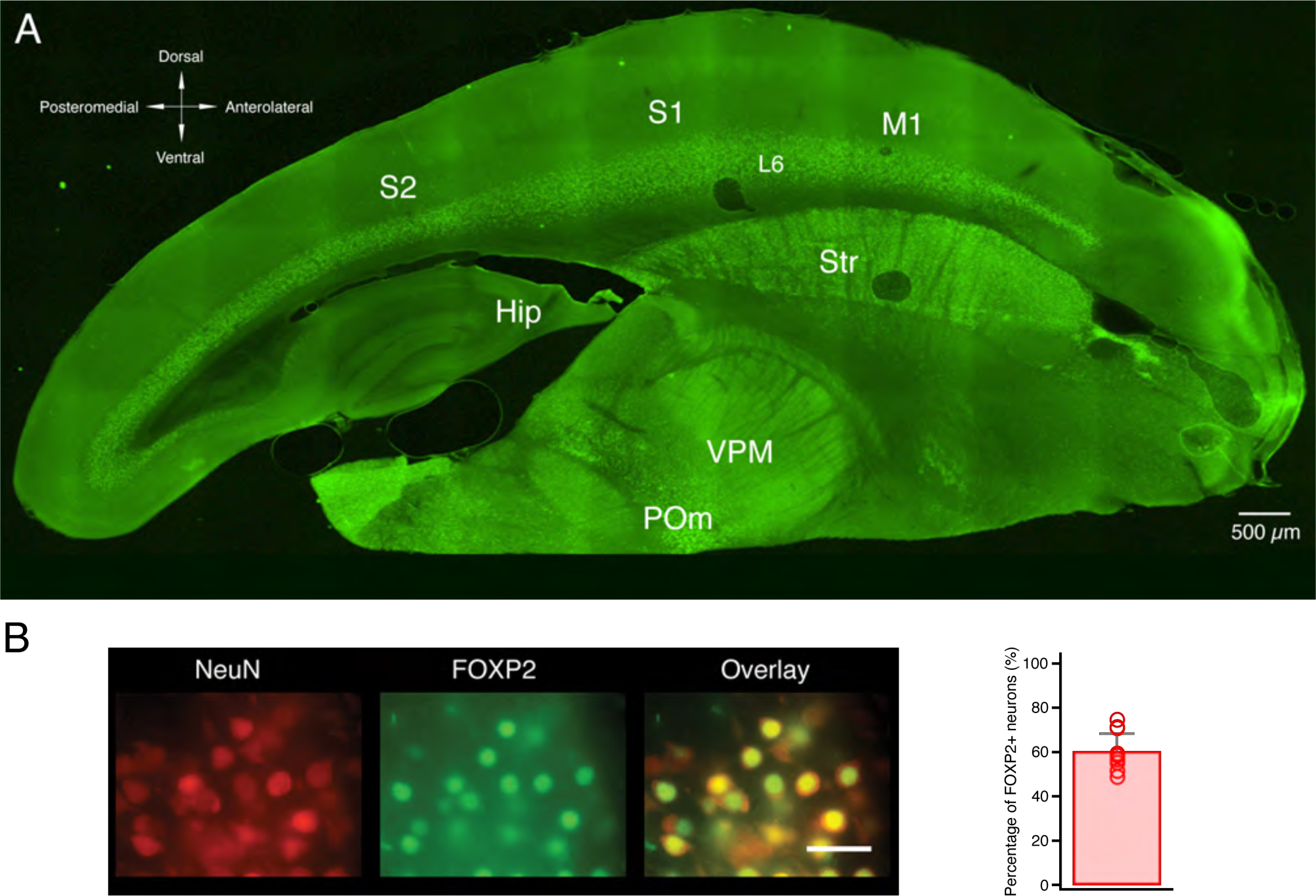
FOXP2 expression in the neocortex. (A) FOXP2 immunofluorescence in a 150 µm thalamocortical rat brain slice (B,C) Co-localisation of NeuN and FOXP2 in L6. High-power micrograph of neurons in L6 (B) that are labelled by NeuN (left) and immunoreactive for FOXP2 (middle); the rightmost images show the relative proportion of FOXP2+ neurons in L6. The bar graph shows the proportion of FOXP2+ neurons relative to the total number of neurons in L6 (n=12). The T-bar represents the standard deviation. In L6, 60% of excitatory neurons are immunoreactive for FOXP2.

### Cell-specific morphological properties of FOXP2+ and FOXP2– excitatory neurons in layer 6a and 6b

To investigate the morphological characteristics of FOXP2+ and FOXP2– neurons in S1 barrel cortex, we performed whole-cell patch-clamp recordings combined with *post hoc* FOXP2 immunolabelling and biocytin staining. In total, 171 L6 excitatory neurons and ten L6 interneurons were recorded, of which 62 excitatory neurons were located in upper layer 6 (layer 6a, defined here as the upper 70% of layer 6) and 97 in layer 6b (**Suppl. Figure S2**); the remaining 12 excitatory neurons were excluded from the analysis because of an ill-defined location (either close to the L5-L6 border or to the L6a-L6b border). After FOXP2 immunolabelling, neurons with high-quality biocytin staining (n=40) were carefully selected for detailed three-dimensional (3D) morphological reconstructions.

Figure 2A,B and **Suppl. Figure S4** show the somatodendritic and axonal domains of representative FOXP2+ and FOXP2– excitatory neurons located in L6a and 6b, respectively. Without exception, FOXP2+ neurons in both layer 6a and 6b were found to be upright PCs with apical dendrites oriented towards to the pial surface. The dendritic domains of L6a and L6b FOXP2+ PCs were confined to their home barrel column and showed no significant differences (**Figure 2**; **Table 1, Suppl. Table S1**). However, the apical dendrite of L6a FOXP2+ PCs predominantly terminated in layer 4, whereas that of L6b FOXP2+ PCs terminated in upper layer 5. Similarly, axon collaterals of L6a and L6b FOXP2+ PCs projected to either layer 4 or upper layer 5, respectively, depending on their location in layer 6 (**Figure 2A,C,E)**. While axon collaterals of L6a FOXP2+ PCs were mainly confined to their home barrel column (Crandall et al., 2017), those of L6b FOXP2+ PCs also innervated adjacent columns (horizontal field-span: 388.5 ± 163.6 µm for L6a FOXP2+ PCs (n=10); 607.1 ± 238.9 µm for L6b FOXP2+ PCs (n=10)) (**Figure 2A,B,E; Suppl. Figure S3; Table 1, Suppl. Table S1**). The primary axons of L6a and L6b FOXP2+ PCs projected deeply into the WM and turned laterally to run parallel to other axons.

**Figure 2.**
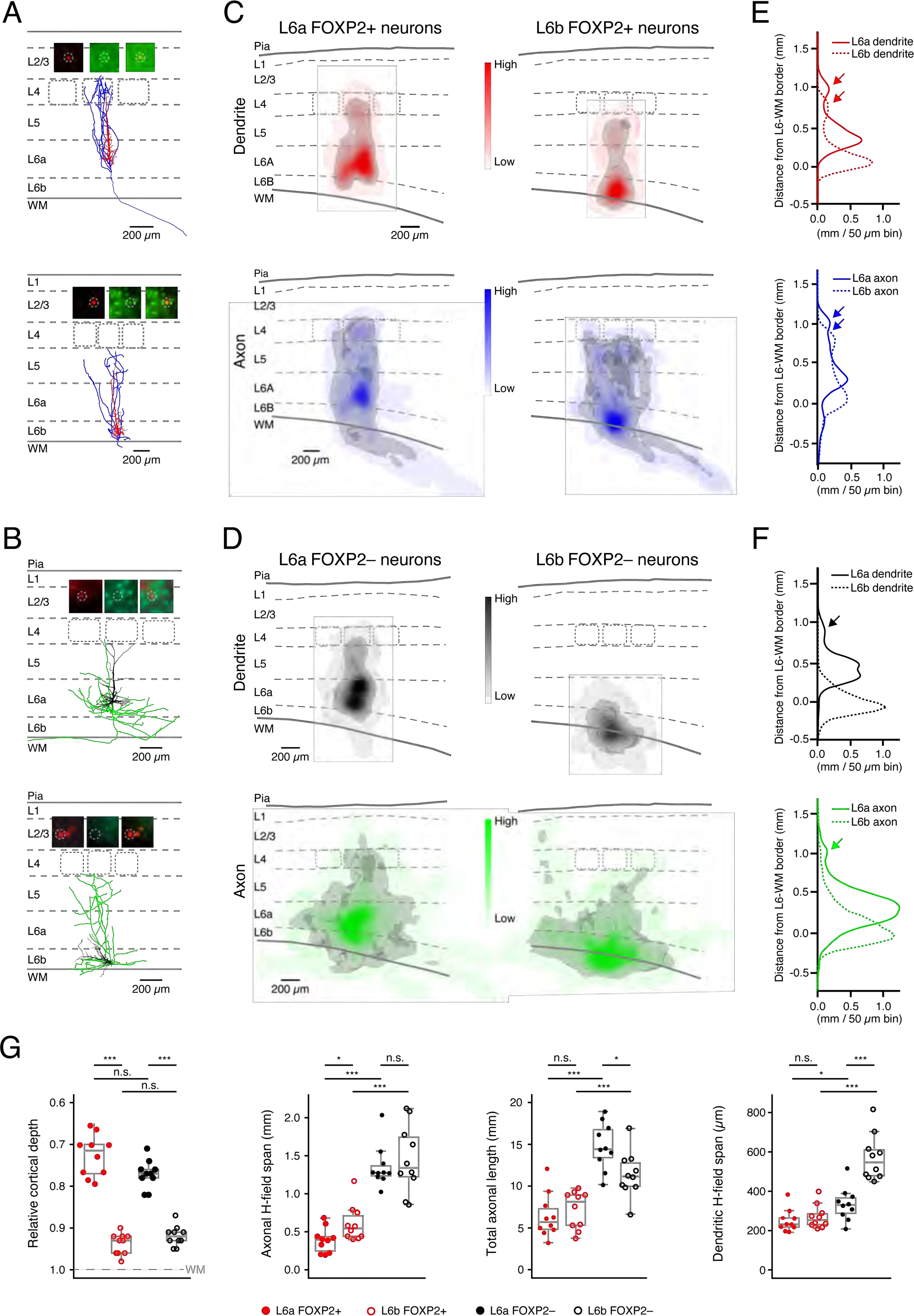
Morphological differences between L6a and L6b FOXP2+ and FOXP2– excitatory neurons. (A) Representative axo-somatodendritic morphology of a L6a FOXP2+ PC (top) and L6b FOXP2+ PC (bottom). Dendrites are in red, axons in blue. Insets show the FOXP2 immunoreactivity (see Methods for details). (B) Representative axo-somatodendritic morphology of a L6b FOXP2– PC (top) and a L6b FOXP2– PC (bottom). Dendrites are in black and axons in green. Insets show the FOXP2 immunoreactivity. (C) 2D projections of 3D density maps for L6a (left) and L6b (right) FOXP2+ PC dendrites and axons; colour code as in (A). The 80-percentile of the integrated dendritic and axonal length density is shown in grey (D) 2D projections of 3D density maps for L6a (left) and L6b (right) FOXP2– neuron dendrites and axons; colour code as in (B). The 80-percentile of the integrated dendritic and axonal length density is shown in grey. (E) 1D dendritic (top, red) and axonal (bottom, blue) densities for L6a and L6b FOXP2+ PCs along the vertical axis. In these plots, 0 mm is the position of the L6/WM border. Solid lines represent L6a neurons, dashed lines L6b neurons. Arrows indicate the dendritic and axonal projections to superficial cortical layers. (F) 1D dendritic (top, black) and axonal (bottom, green) densities for L6a and L6b FOXP2– excitatory neurons along the vertical axis. In these plots, 0 mm is the position of the L6/WM border. Solid lines represent L6a FOXP2– neurons, dashed lines L6b FOXP2– neurons. Note that L6b FOXP2– excitatory neurons comprise a heterogeneous population of non-PC neurons with an dendritic and axonal domain that projects into the WM. Arrows indicate the dendritic and axonal projections to superficial cortical layers. (G) Box plots showing the relative cortical depth, axonal horizontal field span, the total axonal length, and the horizontal dendritic field span for L6a FOXP2+ (n=10), L6b FOXP2+ (n=10), L6a FOXP2– (n=10), and L6b FOXP2– (n=10) excitatory neurons, respectively. The box plot displays the interquartile range (IQR), with whiskers indicating 1.5 times the IQR and a horizontal line marking the median. The statistical significance was assessed using the Wilcoxon Mann-Whitney U test.

**Table 1.** Morphological properties of L6 FOXP2+ and FOXP2– excitatory neurons. For statistical differences between FOXP2+ and FOXP2- neurons in layer 6a and 6b, see Table S1.

|  | **L6a FOXP2+**<br>(n=10) | **L6a FOXP2~**<br>(n=10) | **L6b FOXP2+**<br>(n=10) | **L6b FOXP2–**<br>(n=10) |
| --- | --- | --- | --- | --- |
| ***Soma*** |  |  |  |  |
| Depth (μm) | 1479.0 ± 81.8 | 1384.4 ± 102.0 | 1700.6 ± 155.4 | 1739.6 ± 158.2 |
| Relative depth (%) | 77.7 ± 4.4 | 72.7 ± 5.3 | 91.8 ± 2.5 | 93.7 ± 2.5 |
| perimeter (μm) | 56.7 ± 7.7 | 66.7 ± 11.4 | 54.8 ± 7.6 | 63.5 ± 10.2 |
| area (μm<sup>2</sup>) | 242.6 ± 60.9 | 340.8 ± 110.3 | 229.4 ± 61.3 | 299.3 ± 95.0 |
| ***Dendrite*** |  |  |  |  |
| No. of apical dendrite branches | 30.2 ± 7.2 | 17.6 ± 9.2 | 26.9 ± 8.5 | N/A |
| Total length of apical dendrite (μm) | 3634.2 ± 985.7 | 2485.5 ± 754.3 | 4009.4 ± 714.7 | N/A |
| No. of basal dendrites | 6.8 ± 1.4 | 6.0 ± 1.6 | 7.2 ± 1.5 | 6.0 ± 1.2 |
| No. of basal dendrite branches | 18.7 ± 3.1 | 23.6 ± 8.8 | 20.4 ± 9.3 | 36.3 ± 15.2 |
| Total length of basal dendrites (μm) | 1870.2 ± 391.8 | 3300.2 ± 1026.8 | 2001.2 ± 520.9 | 6222.4 ± 2804.9 |
| Horizontal field span of dendrite (μm) | 249.9 ± 57.0 | 333.9 ± 84.0 | 269.6 ± 59.7 | 570.6 ± 117.9 |
| No. of dendritic branches | 48.9 ± 8.5 | 41.2 ± 15.6 | 47.3 ± 15.6 | 36.3 ± 15.2 |
| Total length of dendrites (μm) | 5504.4 ± 1205.3 | 5785.7 ± 1530.2 | 6010.7 ± 723.0 | 6222.4 ± 2804.9 |
| ***Axon*** |  |  |  |  |
| Horizontal field span of axon (μm) | 388.5 ± 163.6 | 1353.5 ± 274.8 | 607.1 ± 238.9 | 1453.6 ± 445.8 |
| No. of axonal branches | 26.3 ± 11.5 | 49.1 ± 11.9 | 22.2 ± 8.2 | 37.4 ± 9.5 |
| Total length of axon (μm) | 6404.7 ± 2637.6 | 14769.9 ± 2779.7 | 7212.9 ± 2247.9 | 11384.1 ± 2719.5 |

In marked contrast, the dendritic morphology of FOXP2– excitatory neurons was highly heterogeneous both within and between layers (**Figure 2B,D,F,G; Suppl. Figure S3; Table 1**). The majority of L6a FOXP2– excitatory neurons were upright PCs with apical dendrites terminating either in layer 4 or 5. The dendritic arbour of L6a FOXP2– PCs was confined to the home column with a horizontal field-span of 333.9 ± 84.0 µm (n=10). A bipolar or inverted dendritic morphology was observed in a minority of L6a FOXP2– excitatory neurons. Axon collaterals of L6a FOXP2– PCs projected horizontally in infragranular layers with an average field-span of 1353.5 ± 274.8 µm (n=10). Furthermore, tall, putative claustrum-projecting PCs with a wide basal dendritic domain and an apical dendrites terminating in L1 (Cotel et al., 2017; van Aerde and Feldmeyer, 2015; Yang et al., 2022) were also found to be FOXP2–.

In layer 6b, no upright FOXP2– PCs were found. L6b FOXP2– excitatory neurons comprised two main morphological subgroups: inverted pyramidal cells and multipolar spiny neurons (Marx and Feldmeyer, 2013; Zhang and Deschênes, 1997). These neurons have long dendritic and axonal branches with a horizontal field-span of 570.6 ± 117.9 µm (n=10) and 1453.6 ± 445.8 µm (n=10), respectively (**Figure 2B,D,F,G; Suppl. Figure S3; Table 1, Suppl. Table S1**). L6b FOXP2– bipolar and horizontally/tangentially oriented pyramidal cells were only infrequently found. In addition to L6 excitatory neurons, ten L6 interneurons with either fast spiking (n=3) or non-fast spiking (n=7) firing patterns were recorded. All were FOXP2– (**Suppl. Figure S4**) and had diverse axonal domains, ranging from dense local to translaminar/transcolumnar projection patterns.

The average dendritic and axonal arborisation pattern of FOXP2+ and FOXP2– excitatory neurons in layer 6 was determined by calculating 1D and 3D dendritic and axonal density maps (**Figure 2C,D**). Basal dendrites of both L6a and L6b FOXP2+ PCs are mainly located in their respective layers. In addition, local dendritic density maxima were identified in the region where the apical dendrites terminated, i.e. at the L4/5A border and in the middle of L5 for L6a and L6b FOXP2+ CT PCs, respectively (**Figure 2C,D**; see also arrows in **Figure 2E,F**). The L6a FOXP2+ PC axon has only a narrow axonal domain and in S1 barrel cortex innervates only the home barrel column. In contrast, The axon field span of L6b FOXP2+ PCs is larger and innervates also neighbouring barrel columns (**Figure 3C,E**).

**Figure 3.**
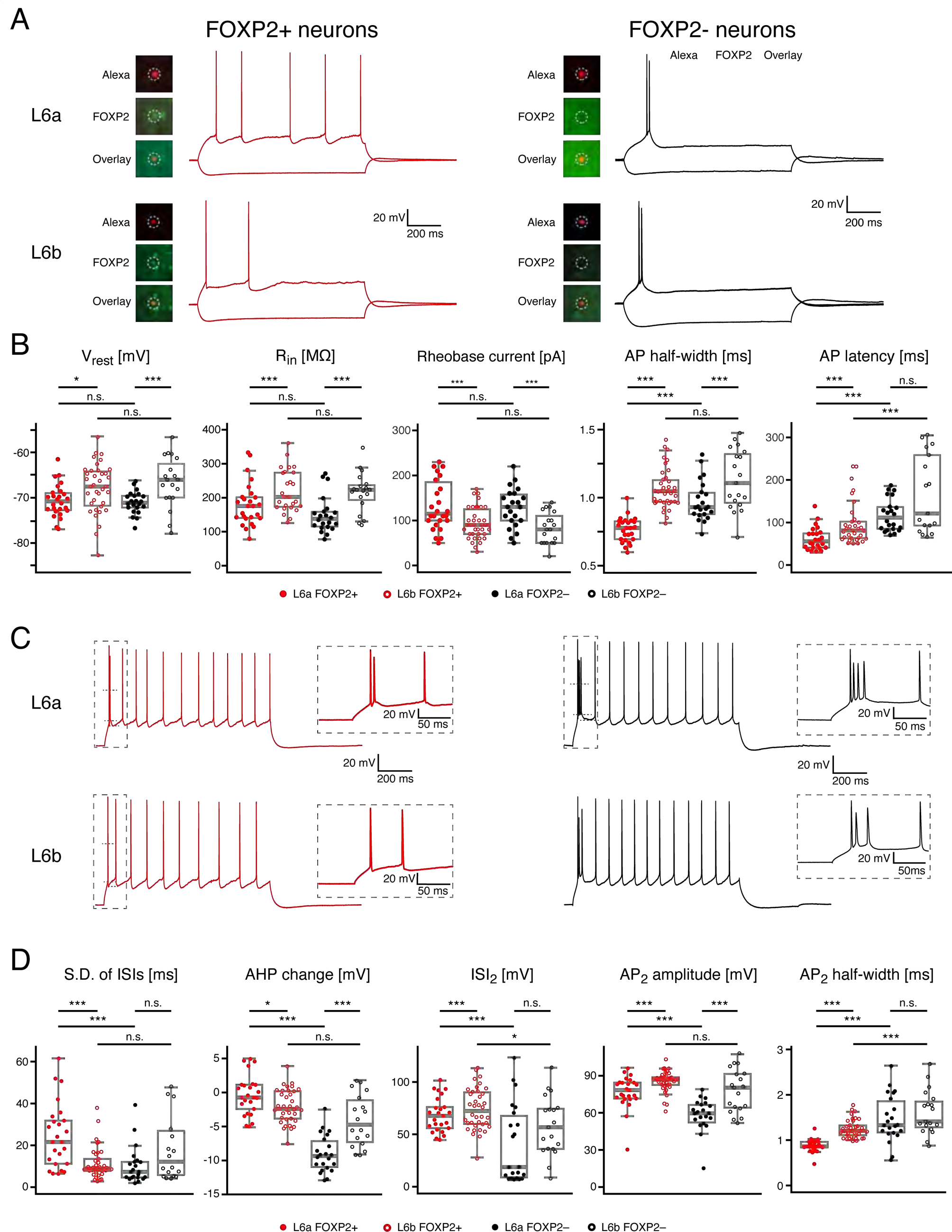
L6a and L6b FOXP2+ and FOXP2– excitatory neurons show distinct electrophysiological properties. (A) Representative recordings of membrane potential changes in response to hyperpolarizing and rheobase current injections for FOXP2+ (left) and FOXP2– (right) excitatory neurons in layer 6a (top) and 6b (bottom), respectively. Colour photographs show Alexa fluorescence labelling of the recorded neuron (red, top), FOXP2 immunoreactivity (green, middle) and the overlay of the fluorescence images (bottom). L6a and L6b FOXP2+ PCs showed only one single AP near the rheobase while FOXP2– excitatory neurons exhibited an initial burst of two to four APs. (B) Box plots for passive electrophysiological properties and of the first AP of L6a FOXP2+ (n=26, red closed circles), L6b FOXP2+ (n=22, red open circles), L6a FOXP2– (n=38; black closed circles) and L6b FOXP2– (n=19, black open circles) excitatory neurons. (C) Representative recordings of repetitive firing patterns of L6a nd L6b FOXP2+ (left) and FOXP2– (right) excitatory neurons. The insets in dashed boxes show an initial spike doublet (L6a and L6b CT PC) or AP burst (L6a and L6b CC neuron) at higher temporal resolution. (D) Box plots for repetitive firing properties of L6a FOXP2+, L6b FOXP2+, L6a FOXP2–, and L6b FOXP2– excitatory neurons (labelling as in panel B). Boxes indicates the interquartile range (IQR), the whiskers show the range of values that are within 1.5_*_IQR and a horizontal line indicates the median. The statistical significance was assessed using the Wilcoxon Mann-Whitney U test.

L6a FOXP2+ and FOXP2– PCs have largely similar dendritic domains. In contrast, the dendritic domain of L6b FOXP2– excitatory neurons is markedly different from that of L6b FOXP2+ PCs, being almost exclusively located close to the L6b-WM border (**Figure 2D,F**). The axonal density profiles of both L6a and L6b FOXP2– excitatory neurons span several barrel columns in the horizontal direction, indicating a corticocortical axonal projection. Axons of L6a FOXP2– PCs innervate both granular and infragranular layers, whereas axons of L6b FOXP2– excitatory neurons project mainly within L6 and the WM.

### Cell-specific electrophysiological properties of FOXP2+ and FOXP2– excitatory neurons in layer 6a and 6b

The intrinsic membrane properties of L6a FOXP2+ (n=26), L6b FOXP2+ (n=22), L6a FOXP2– (n=38) and L6b FOXP2– excitatory neurons (n=19) including the passive electrophysiological properties and AP characteristics (**Figure 3**) were analysed. L6a and L6b FOXP2+ excitatory neurons were found to be significantly different from L6a and L6b FOXP2– excitatory neurons in almost all electrophysiological properties (**Figure 3**, **Table 2, Suppl. Table S2**). The rheobase current was significantly larger for L6a than for L6b FOXP2+ and FOXP2– PCs. At near rheobase current injection, L6a and L6b FOXP2+ CT PCs generated a single AP that was sometimes followed by a short AP train. Under the same condition, L6a and L6b FOXP2– CC PCs produced a spike doublet or a short AP burst followed by a prominent depolarising afterpotential (DAP; **Figure 3A**). The AP onset latency of L6a and L6b FOXP2+ excitatory neurons was shorter than in L6a and L6b FOXP2– excitatory neurons, respectively. The AP half-width and latency was shortest in L6a FOXP2+ excitatory neurons while the AP threshold and AP amplitude did not differ significantly between the four excitatory neuron groups (**Figure 3B**; **Table 2**).

**Table 2.** Electrophysiological properties of L6 FOXP2+ and FOXP2– excitatory neurons. For statistical differences between FOXP2+ and FOXP2- neurons in layer 6a and 6b, see Table S2.

|  | **L6a FOXP2+**<br>(n=26) | **L6b FOXP2+**<br>(n=8) | **L6a FOXP2–**<br>(n=22) | **L6b FOXP2–**<br>(n=19) |
| --- | --- | --- | --- | --- |
| ***Passive*** |  |  |  |  |
| V<sub>rest</sub> (mV) | -70.6 ± 3.6 | -68.3 ± 6.3 | -71.1 ± 2.5 | -66.6 ± 5.6 |
| R<sub>in</sub> (MΩ) | 180.7 ± 63.5 | 222.0 ± 57.0 | 149.5 ± 53.5 | 216.0 ± 58.2 |
| τ<sub>m</sub> (ms) | 16.2 ± 4.4 | 20.6 ± 5.2 | 19.6 ± 4.2 | 23.7 ± 7.0 |
| Sag (%) | 12.9 ± 7.7 | 13.3 ± 7.1 | 10.6 ± 6.3 | 5.6 ± 4.4 |
| ***Single AP*** |  |  |  |  |
| Rheobase current (pA) | 141.5 ± 76.4 | 93.7 ± 34.8 | 127.7 ± 43.7 | 83.4 ± 34.6 |
| AP threshold (mV) | -36.4 ± 2.8 | -37.5 ± 4.1 | -37.5 ± 2.9 | -37.6 ± 5.7 |
| AP half-width (ms) | 0.76 ± 0.09 | 1.07 ± 0.14 | 0.97 ± 0.15 | 1.13 ± 0.22 |
| AP amplitude (mV) | 91.9 ± 6.4 | 95.2 ± 5.7 | 90.3 ± 8.2 | 94.4 ± 10.2 |
| AP latency (ms) | 62.1 ± 27.0 | 97.6 ± 50.8 | 117.2 ± 37.9 | 166.9 ± 91.6 |
| fAHP amplitude (mV) | 10.1 ± 2.6 | 11.4 ± 4.5 | 3.4 ± 2.6 | 9.2 ± 5.7 |
| fAHP latency (ms) | 3.4 ± 2.6 | 27.2 ± 29.9 | 20.0 ± 36.2 | 16.5 ± 21.3 |
| ***Repetitive firing*** |  |  |  |  |
| Max. firing frequency (Hz) | 24.8 ± 7.1 | 21.4 ± 5.6 | 23.5 ± 8.6 | 20.6 ± 4.4 |
| Slope of F-I curve (APs/100 pA) | 11.8 ± 4.1 | 15.4 ± 6.2 | 12.1 ± 4.5 | 15.5 ± 7.9 |
| Adaptation ratio | 1.1 ± 0.3 | 1.1 ± 0.2 | 1.1 ± 0.2 | 1.4 ± 0.5 |
| s.d. of ISIs (ms) | 24.6 ± 15.3 | 11.5 ± 7.6 | 10.2 ± 8.9 | 20.6 ± 20.1 |
| ISI1 (ms) | 19.2 ± 24.2 | 36.1 ± 20.7 | 6.7 ± 2.7 | 23.1 ± 15.4 |
| ISI2 (ms) | 68.6 ± 16.4 | 74.2 ± 19.9 | 40.6 ± 37.1 | 58.0 ± 28.1 |
| ISI3 (ms) | 87.7 ± 27.7 | 85.4 ± 14.1 | 83.5 ± 38.1 | 72.3 ± 22.0 |
| AP1 amplitude (mV) | 92.7 ± 5.7 | 94.8 ± 5.9 | 91.1 ± 5.7 | 92.8 ± 9.2 |
| AP2 amplitude (mV) | 76.7 ± 13.2 | 85.3 ± 8.3 | 56.4 ± 16.3 | 77.9 ± 17.4 |
| AP3 amplitude (mV) | 82.1 ± 9.2 | 86.2 ± 6.9 | 72.3 ± 14.5 | 84.3 ± 11.6 |
| AP1 half-width (ms) | 0.73 ± 0.08 | 1.00 ± 0.14 | 0.92 ± 0.11 | 1.05 ± 0.22 |
| AP2 half-width (ms) | 0.88 ± 0.14 | 1.25 ± 0.20 | 1.47 ± 0.52 | 1.60 ± 0.48 |
| AP3 half-width (ms) | 0.88 ± 0.12 | 1.24 ± 0.20 | 1.46 ± 0.38 | 1.51 ± 0.40 |
| AP1 threshold (mV) | -40.1 ± 3.6 | -39.2 ± 4.2 | -38.6 ± 3.6 | -38.2 ± 5.7 |
| fAHP1 amplitude (mV) | 6.3 ± 3.4 | 5.3 ± 3.5 | 0.0 ± 2.7 | 3.9 ± 4.6 |
| AP9 threshold (mV) | -32.8 ± 4.4 | -34.0 ± 4.9 | -34.1 ± 4.4 | -34.4 ± 6.7 |
| fAHP9 amplitude (mV) | 14.1 ± 2.0 | 12.5 ± 2.4 | 13.5 ± 1.7 | 11.8 ± 2.6 |
| fAHP9 - fAHP1 (mV) | -0.4 ± 2.9 | -2.1 ± 2.4 | -9.0 ± 2.7 | -4.1 ± 3.8 |

Repetitive firing properties of FOXP2+ and FOXP2– excitatory neurons in layer 6a and 6b were analysed using a current injection, which elicited ∼10 APs. In L6a and L6b FOXP2+ neurons a single AP or a spike doublet occurred at the beginning of the AP train and was followed by irregular spiking, with variable inter-spike intervals (ISIs) (**Figure 3C**). In contrast, L6a and L6b FOXP2– excitatory neurons showed an initial burst of up to four APs, followed by regular spiking with constant ISIs. Furthermore, in FOXP2+ excitatory neurons, the AHP amplitude remained constant, whereas in FOXP2– excitatory neurons, AHPs became progressively larger during the spike train (**Figure 3C**). Furthermore, significant differences were observed in the duration of the 2^nd^ ISI, the amplitude and half-width of 2^nd^ AP in a spike train between FOXP2+ and FOXP2– excitatory neurons in both layer 6a and 6b (**Figure 3D**, **Table 2**).

L6b FOXP2+ and FOXP2– excitatory neurons exhibited a largely similar dichotomy in firing patterns as L6a FOXP2+ and FOXP2– excitatory neurons, with the exception of a small subset of L6b FOXP2– excitatory neurons (see below). Specifically, L6b FOXP2+ excitatory neurons also displayed only an initial single AP or a AP doublet followed by an irregular AP firing with variable ISIs. In contrast, L6b FOXP2– excitatory neurons showed an initial burst of up to four APs with a subsequent regular spiking with constant ISIs (**Suppl. Figure S5**). Of those, the majority of L6b FOXP2– excitatory neurons displayed a ‘rapidly adapting’ firing pattern, with a long ISI between the initial burst and following APs but shorter, stable ISIs thereafter. L6b FOXP2– excitatory neurons with a rapidly adapting firing patterns exhibit an inverted pyramidal morphology. Another subpopulation of L6b FOXP2– excitatory neurons showed no initial AP burst but a slowly adapting firing pattern, with increasingly longer ISIs. These neurons had a multipolar dendritic morphology (**Suppl. Figure S5**). A thorough examination of the intrinsic membrane properties of the two populations of L6b FOXP2– excitatory neurons showed differences in input resistance (R_in_), 1^st^ AP latency, adaptation ratio, s.d. of ISIs, AHP change and several other parameters (**Suppl. Figure S6** and **Table S1**).

In summary, L6a and L6b FOXP2+ PCs exhibit differences in several electrophysiological properties (e.g. R_in_, AP half-width, s.d. of ISIs, see **Figure 3** and **Table 2**) and their dendritic and axonal arborisation pattern (Figure 2 and **Table 1**). FOXP2– excitatory neurons in layer 6 comprise three morpho-electrophysiological phenotypes: L6a upright PCs with rapidly adapting (or burst spiking) firing patterns, L6a and L6b inverted PCs with rapidly adapting firing patterns, and L6b multipolar excitatory neurons comprising neurons with no apparent main dendrite or a tangential/ oblique main dendrite (Marx and Feldmeyer, 2013) and a slowly adapting firing pattern (**Figure 2**, **Figure 3**, **Table 1**, **Table 2, Suppl. Figure S5**).

### Synaptic connections established by L6a and L6b FOXP2+ and FOXP2– excitatory neurons have distinct functional properties

Paired recordings from synaptically coupled L6 FOXP2+ and FOXP2– excitatory neurons were conducted to reveal potential differences in EPSP time course, neurotransmitter release probability, and synaptic dynamics. Following the electrophysiological recordings, FOXP2 antibody labelling and subsequent biocytin staining was performed. A total of 24 synaptic connections were obtained. Nine were established by presynaptic L6a FOXP2+ PCs, three by L6b FOXP2+ PCs, eight by L6a FOXP2– PCs, and four by L6b FOXP2– excitatory neurons. Postsynaptic neurons in these connections include both L6a and L6b FOXP2+ and FOXP2– excitatory neurons as well as L6b fast spiking (FS) and non-fast spiking (nFS) interneurons. All connections with a presynaptic FOXP2+ neuron were weak (1^st^ EPSP amplitude for presynaptic L6a FOXP2+ PCs: 0.13 ± 0.13 mV; for presynaptic L6b FOXP2+ PCs: 0.29±0.24 mV) and displayed strong short-term facilitation (paired-pulse ratio EPSP_2_/EPSP_1_ for presynaptic L6a FOXP2+ PCs: 2.78±1.65; for presynaptic L6b FOXP2+ PCs: 3.38±3.80) (**Figure 4**). This suggests that the initial synaptic release probability at this connection types is low. On the other hand, connections with a presynaptic FOXP2– neuron were comparatively strong (1^st^ EPSP amplitude for presynaptic L6a FOXP2– PCs: 0.57±0.31 mV; for presynaptic L6b FOXP2– PCs: 0.35±0.17 mV) and showed short-term depression with a paired-pulse ratio of (paired-pulse ratio EPSP_2_/EPSP_1_ for presynaptic L6a FOXP2– PCs: 0.86±0.11; for presynaptic L6b FOXP2– PCs: 0.82±0.22) (**Figure 4**). Thus, synaptic connections established by presynaptic L6 FOXP2+ and FOXP2– excitatory neurons demonstrate a highly distinctive synaptic efficacy and short-term plasticity, indicating presynaptic cell-type specificity.

**Figure 4.**
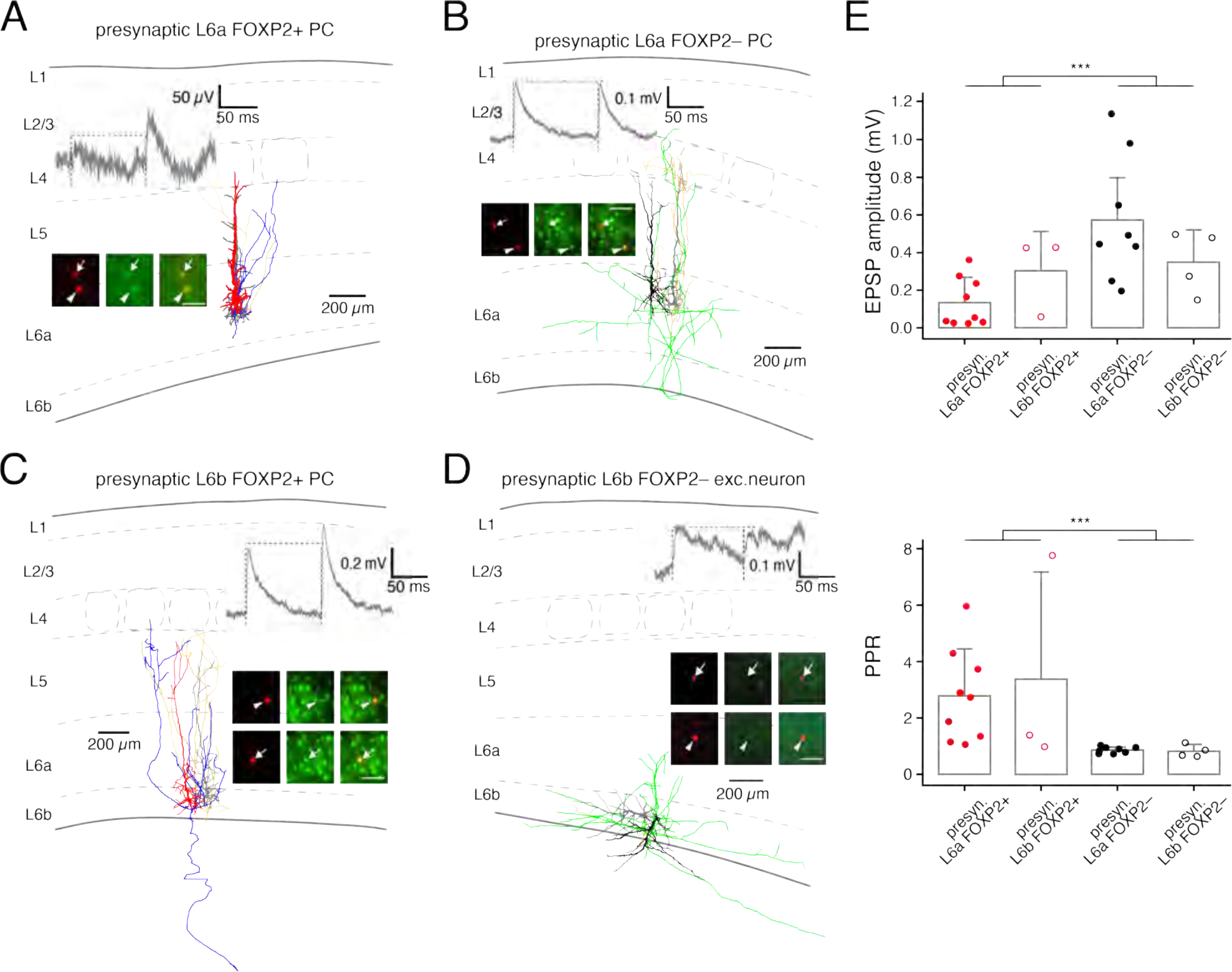
Synaptic connections established by L6 FOXP2+ and FOXP2– excitatory neurons show distinct properties. (A-D) L6a and L6b synaptic connections; insets, FOXP2 immunoreactivity. Morphology, electrophysiology and FOXP2 immunoreactivity for synaptic connections between (A) two L6a FOXP2+ CT PCs, (B) a L6a FOXP2– CC and a L6a FOXP2+ CT PC, (C) two L6b FOXP2+ CT PCs, and (D) two L6b FOXP2– inverted PCs. Colour code: red dendrite/blue axon: L6a and L6b FOXP2+ CT PC, black dendrite/green axon: L6a and L6b FOXP2– CC excitatory neuron, grey dendrite/dark yellow axon: postsynaptic L6a and L6b excitatory neuron. (E) Histogram for the 1^st^ EPSP amplitude (top) and paired-pulse ratio (bottom) of synaptic connections established by L6a FOXP2+ (n=9), L6b FOXP2+ (n=3), L6a FOXP2– (n=8), and L6b FOXP2– (n=4) neurons, respectively. Synaptic connections for which the presynaptic neuron was identified only on morphological grounds (axonal domain, cortical depth, and electrophysiological properties) have been included in the analysis. The box indicates the mean and the whiskers show the s.d. P values were calculated using the Wilcoxon Mann-Whitney U test.

### FOXP2-immunopositive neurons in layer 6 are corticothalamic neurons

Layer 6 is the source of a high proportion of corticothalamic afferents (Bourassa et al., 1995; Zhang and Deschênes, 1997). We investigated whether corticothalamic projections originated from FOXP2+ or FOXP2– PCs. To achieve this, choleratoxin subunit B (CTB) conjugated with Alexa Fluor 488 and 647 was injected in the same animal into the VPM and POm, respectively (**Figure 5A**).

**Figure 5.**
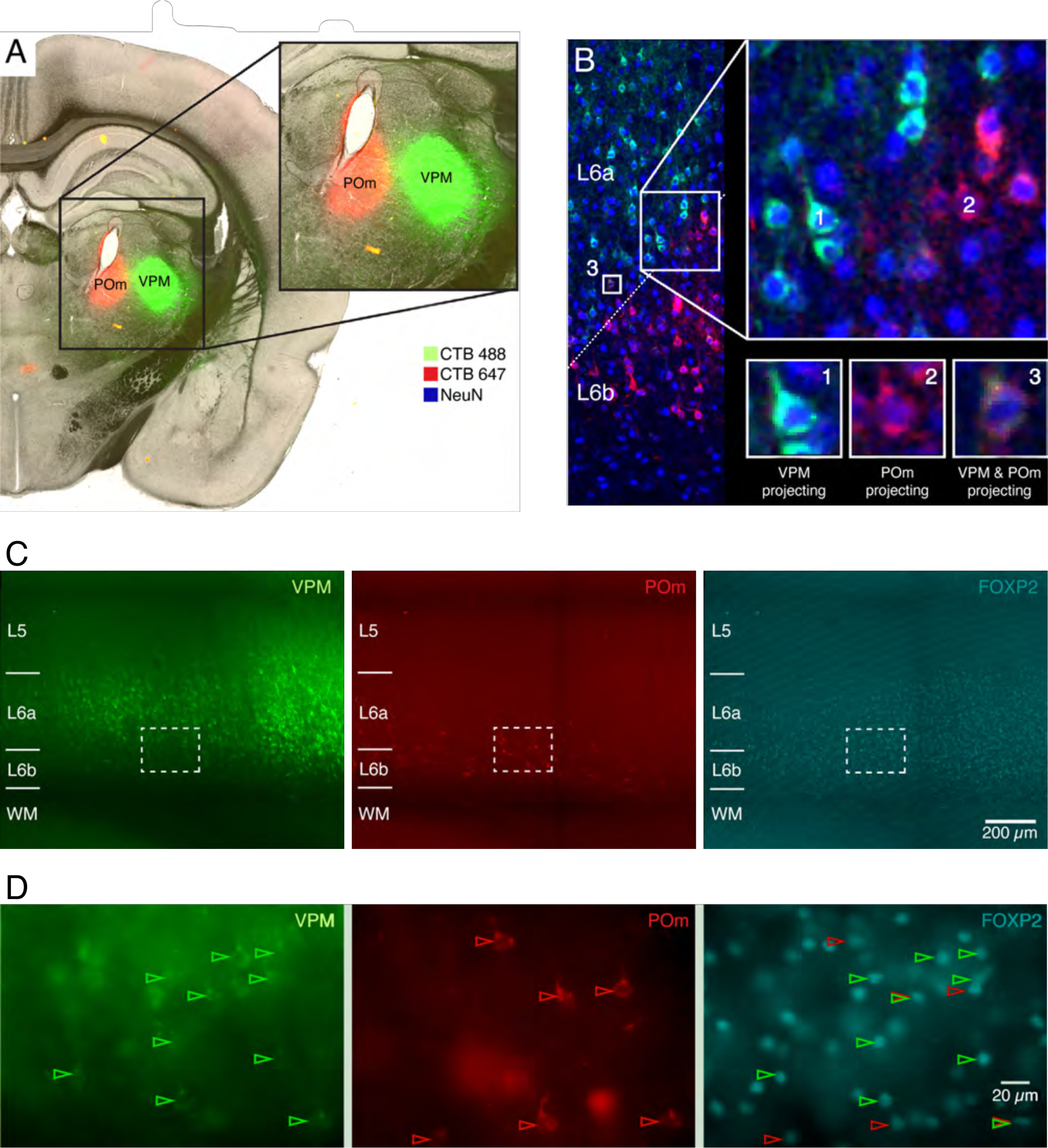
FOXP2–positive neurons are corticothalamic (CT) neurons that project to the VPM or POm or both. (A) Cholera toxin subunit B (CTb) conjugated with two Alexa Fluor dyes, A488 and A647, was separately microinjected into the VPM and POm thalamic nuclei. (B) Retrogradely labelled neurons project to either the VPM exclusively (‘green label’, 1), the POm exclusively (‘red label’, 2), or both the VPM and POm (3). NeuN fluorescence labelling is in blue. The oblique dashed line in the leftmost panel marks the border region between layer 6a and 6b. Note the projection target specificity of L6a and L6b CT neurons. (C) Neurons labelled retrogradely via the VPM or both the VPM and POm are located in the superficial and middle layers of layer 6, respectively, as indicated by the FOXP2 labelling (on the right). The majority of L6 PCs which project exclusively to POm is situated in deep layer 6/layer 6b. (D) The areas enclosed by squares in (C) are shown as enlarged images. Green arrows mark L6 neurons projecting to VPM and red arrows those projecting to POm; overlay of red and green arrows indicate neurons projecting to both thalamic nuclei.

The majority of neurons in layer 6, identified by the neuron-specific NeuN stain (**Figure 5B**), were retrogradely labelled by only one of the injected CTBs. Retrogradely labelled neurons projecting to the VPM were predominantly located in layer 6a (**Figure 5C**), whereas retrogradely labelled neurons projecting to the POm were predominantly located in layer 6b (**Figure 5C**). In the transition zone between layer 6a and 6b, a subpopulation of double retrogradely labelled neurons was observed, indicating that they innervate both the VPM and POm (**Figure 5B**). These neurons intermingled with those sending axonal projections to only one thalamic nucleus. All retrogradely labelled neurons in layer 6 were found to be FOXP2+, irrespective of whether they projected to the VPM, POm or both (**Figure 5C, D**). In contrast, POm-projecting, retrogradely labelled neurons in layer 5 exhibited no FOXP2 immunoreactivity (**Suppl. Figure S6**). Thus, FOXP2+ neurons represent all subpopulations of CT neurons in layers 6a and 6b, but not layer 5.

### Differential cholinergic modulation of L6 FOXP2+ and FOXP2– excitatory neurons

L6a and L6b CT PCs have been shown to express both nicotinic and muscarinic acetylcholine receptors (nAChRs and mAChRs; Sorensen et al., 2015; Yang et al., 2020). The effect of 30 µM ACh was tested in 81 L6 excitatory neurons located in both sublayers. In the presence of 0.5 µM TTX to block AP firing, ACh evoked a depolarisation of 10.1 ± 6.6 mV (n=21) in L6a CT PCs and of 18.9 ± 8.4 mV (n=29) in L6b CT PCs (Figure 6A,B), with the ACh-induced depolarisation being significantly larger in L6b than in L6a CT PCs (p < 0.001). In contrast, the ACh response showed greater variability in L6a and L6b FOXP2– neurons. In L6a CC FOXP2– excitatory neurons, 30 µM ACh resulted in either a hyperpolarisation (10 out of 12; ΔV_m_: –1.4±0.8 mV) or a weak depolarisation (ΔV_m_: 1.2 and 2.1 mV, n=2). The majority of L6b CC (FOXP2–) neurons showed an ACh-induced depolarisation (6.3±4.5 mV, n=16 out of 19); in the remaining three, ACh caused a hyperpolarisation (2.1±1.0 mV; **Figure 6A,B**). In the absence of TTX, 30 µM ACh evoked AP firing in 62% of all tested L6a (FOXP2+) CT PCs, 77% L6b (FOXP2+) CT PCs and 45% of L6b (FOXP2–) CC excitatory neurons but not in any L6a (FOXP2–) CC PCs (**Figure 6C**). Thus, L6 CT FOXP2+ PCs are extremely responsive to ACh; in addition L6b CC FOXP2– neurons show also a strong response to ACh.

**Figure 6.**
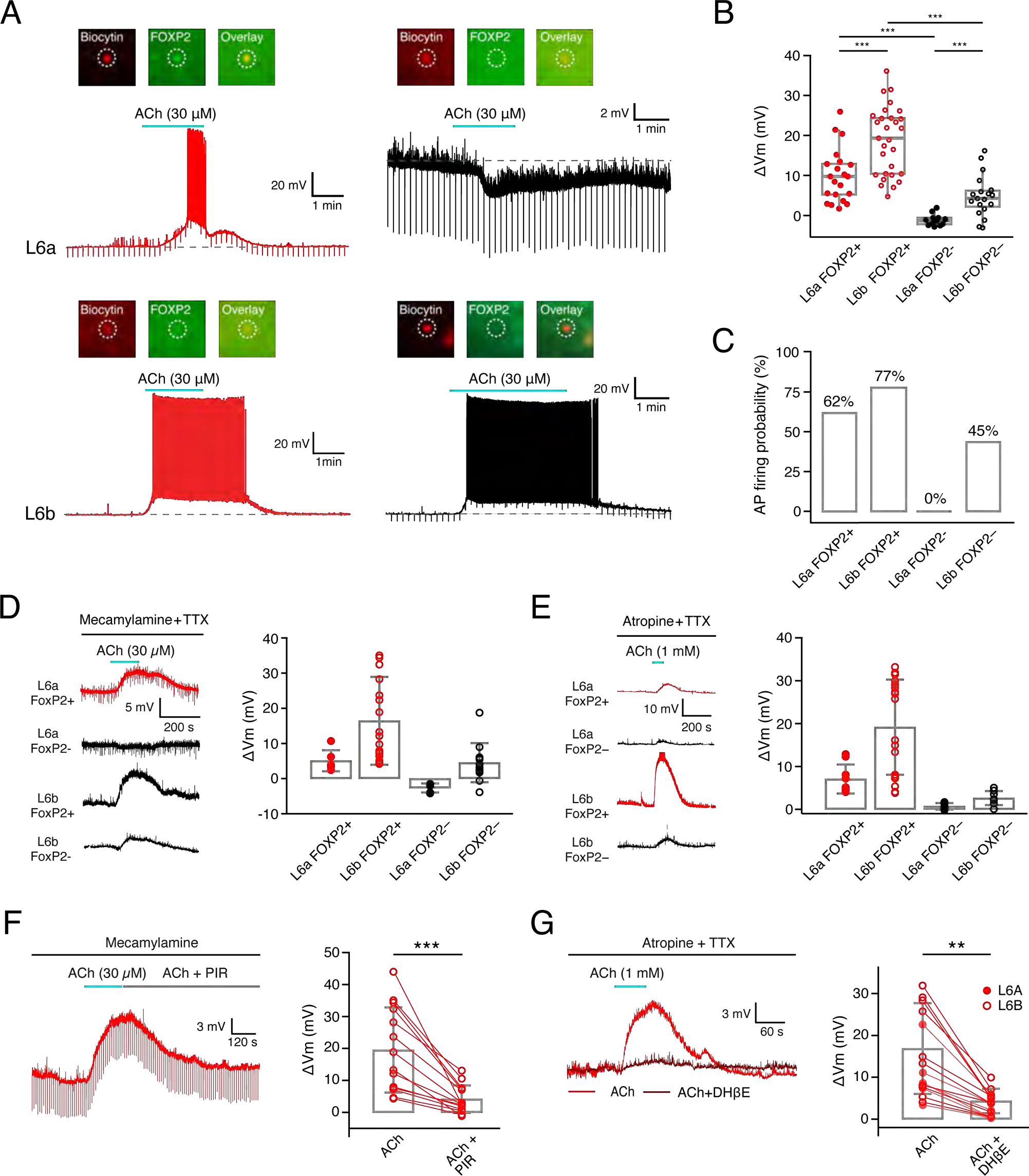
Cholinergic neuromodulation of L6a and L6b FOXP2+ and FOXP2– CT and CC excitatory neurons. (A) Representative voltage responses of L6a (top row) and L6b (bottom row) FOXP2+ and FOXP2– excitatory neurons, respectively, to bath-application of ACh (30 µM). The insets at the top show the immunoreactivity of FOXP2. (B) Box plots for ACh-induced membrane potential changes in L6a FOXP+ CT (n=21), L6a FOXP– CC PCs (n=12), and L6b FOXP– CC excitatory neurons (n=19) in the presence of 0.5 µM TTX to block AP firing. The ACh response of all four L6 excitatory neuron types was significantly different from one another. (C) Fraction of different L6 excitatory neuron types firing action potential following application of 30 µM ACh in the absence of TTX. Note that L6a and L6b FOXP2+ CT PCs but also L6b FOXP2– CC excitatory neurons are highly responsive to ACh and show prolonged trains of spike trains. (D) The muscarinic component of the ACh response in L6 excitatory neurons was isolated by applying 30 µM ACh in the continuous presence of 1 µM mecamylamine and 0.5 µM TTX. L6a and L6b FOXP2+ CT PCs as well as L6b excitatory neurons showed an ACh-induced depolarisation whereas L6a FOXP2– CC excitatory neurons exhibited a hyperpolarising response. Left, original recording of ACh-induced depolarisations in L6a and L6b FOXP2+ CT PCs and FOXP2– CC excitatory neurons; right, bar graphs showing the amplitude of the depolarisation for the four different cell types. (E) The nicotinic component of the ACh response in L6 excitatory neurons was isolated using 200 nM atropine and 0.5 µM TTX. In L6 FOXP2+ CT PCs 1 mM ACh elicited a large nAChR response while in L6a and L6b FOXP2– CC excitatory neurons no or only a small response, respectively, was observed. Left, original recordings of ACh-induced depolarisations in L6a and L6b FOXP2+ and FOXP2– (CT and CC) excitatory neurons; right, bar graphs showing the amplitude of the nAChR-induced depolarisation for the four different cell types. (F) Left, response of a L6 FOXP2+ CT PC to application of 30 µM ACh in the presence of 1 µM mecamylamine, a general nAChR antagonist. This ACh-induced depolarisation was blocked by 0.5 µM pirenzipine, a M_1_ mAChR antagonist. Right, bar graphs show that pirenzipine blocked or largely reduced the mAChR-related depolarisation induced by 30 µM ACh in recorded L6 FOXP2+ CT PCs. (G) Left, response of a L6 FOXP2+ CT PC following application of 1 mM ACh in the presence of the general mAChR antagonist atropine (200 nM) and TTX (0.5 µM). The response was largely blocked by 10 µM DHβE, a specific antagonist of α_4_β_2_* subunit-containing nAChRs. Right, bar graphs show that DHβE blocked or largely reduced the nAChR-related depolarisation induced by 1 mM ACh in recorded L6 FOXP2+ CT PCs. For statistical comparison of the data shown in (B), (D) and (E) the Wilcoxon Mann-Whitney U test was used and the data shown in (F) and (G) the Wilcoxon signed-rank test was used.

To isolate the muscarinic component of the ACh response in L6 FOXP2+ and FOXP2– excitatory neurons, 30 µM ACh was applied in the presence of 0.5 µM TTX and 1 µM mecamylamine, a general nAChR antagonist. As illustrated in **Figure 6D**, L6a and L6b FOXP2+ CT PCs showed a depolarisation of 5.1±3.0 mV (n=6) and 16.4±12.5 mV (n=20), respectively, following ACh application. Conversely, L6a FOXP2– CC excitatory neurons showed a hyperpolarising response to ACh (−2.7±1.3 mV, n=5), consistent with previous findings (Yang et al., 2020) while L6b FOXP2– CC neurons exhibited a small depolarisation (4.5±5.6 mV, n=12).

To determine the contribution of nAChRs to the ACh-induced response in L6a and L6b excitatory neurons, 1 mM ACh was bath-applied together with 200 nM atropine, a general mAChR antagonist, and 0.5 µM TTX. This resulted in a nAChR-mediated depolarisation of 7.0±3.4 mV and 19.1±11.1 mV in L6a and L6b FOXP2+ CT PCs, respectively. On the other hand, the nAChR response in L6a and L6b FOXP2– CC neurons was comparatively small (0.7±0.8 mV and 2.6±1.6 mV, respectively; see **Figure 6E**).

Antagonists for mAChR and nAChR subtypes were used to determine the specific receptor subtypes that contribute to the cholinergic response in L6a and L6b FOXP2+ PCs (**Figure 6F,G**). The muscarinic component was determined by applying 30 µM ACh, 1 µM mecamylamine and 0.5 µM TTX. Subsequent co-application of ACh and 1 µM pirenzipine, an M_1_ mAChR antagonist caused a reduction of the mAChR-mediated depolarisation, indicating that this mAChR subtype mediates the muscarinic component of the ACh response (**Figure 6F**). As shown previously, the hyperpolarising response observed in L6a CC neurons was mediated by M_4_ mAChRs (Yang et al., 2020).

Because the nAChR-mediated response was negligibly small in L6 FOXP2– CC neurons, the nAChR subtype was only determined for L6 CT PCs. For this, ACh was applied in the presence of the specific antagonist for α_4_β_2_* subunit-containing nAChR DHβE (1 µM). In L6a and L6b FOXP2+ CT PCs, DHβE reduced the nAChR response from 10.0±6.0 mV to 1.9±2.4 mV and from 16.9±10.9 mV to 4.3±2.9 mV, respectively, suggesting that it was largely mediated by α_4_β_2_* nAChRs (**Figure 6G**). The data indicate that L6b FOXP2+ CT PCs are particularly responsive to ACh through activation of both M_1_ mAChRs and α_4_β_2_* nAChRs.

ACh caused an increase of neurotransmitter release at synapses established by L6a CT PCs but a decrease at L6a CC PCs (Yang et al., 2020); however, its effect at L6b excitatory synapses has not been characterised until now. We tested how 30 µM ACh affected synaptic transmission in four L6b excitatory neuron pairs (two pairs between L6b FOXP2+ CT PCs and two between L6b FOXP2– inverted PCs; **Suppl. Figure S7**). At the two L6b CT connections ACh caused a marked increase in the EPSP amplitude (from 14 to 99 µV and from 0.42 to 0.58 mV) and a decrease in the paired-pulse ratio (from 6.3 to 0.4 and 1.0 to 0.5). In contrast, the EPSP amplitude at the two L6b inverted PC connections decreased in the presence of ACh (from 0.27 to 0.10 mV and 0.15 to 0.07 mV, respectively) and the paired-pulse ratio increased (from 1.1 to 2.2 and 0.6 to 2.2). Thus, as found for L6a PC connections, ACh selectively enhances the synaptic output of L6b CT PCs and reduced that of L6b CC neurons.

### Differential dopaminergic modulation of L6 FOXP2+ and FOXP2– excitatory neurons

The expression of the *Drd1* gene has been used as a marker for L6b CT PCs (Ansorge et al., 2020; Gong et al., 2003; Hoerder-Suabedissen et al., 2018; Zolnik et al., 2020, Zolnik et al., 2023) but so far dopaminergic responses from L6 CT and CC excitatory neurons have not been studied. Therefore, we examined whether L6a and L6b excitatory neurons are differentially modulated by dopamine (**Figure 7**).

**Figure 7.**
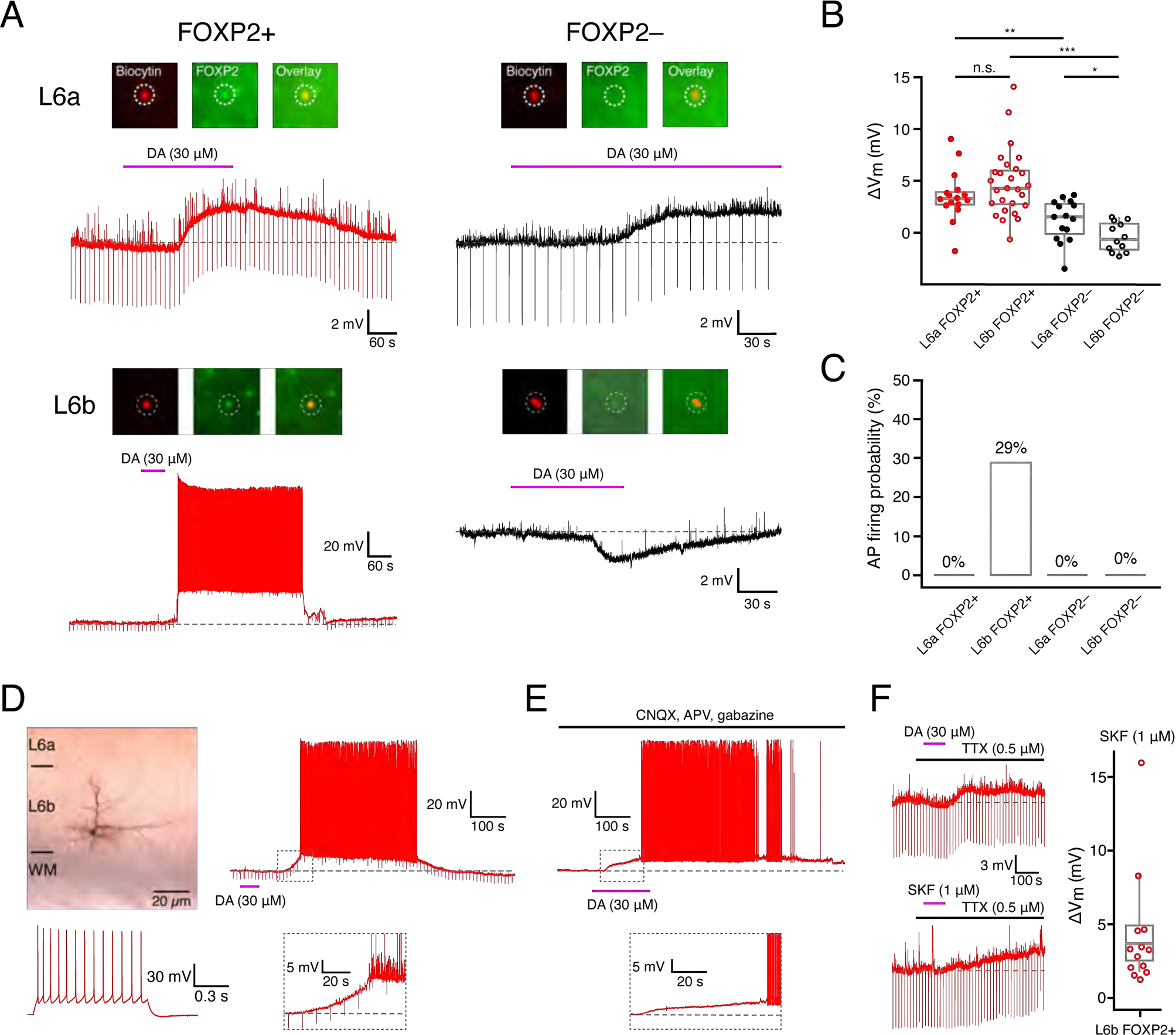
Neuromodulation of L6a and L6b FOXP2+ CT and FOXP2– CC excitatory neurons by dopamine. (A) Representative voltage responses of L6a (top row) and L6b FOXP2+ and FOXP2– excitatory neurons (bottom row), respectively, to bath-application of 30 µM dopamine (DA). The inset at the top show the FOXP2 immunoreactivity of the recorded neurons. (B) Box plots for DA-induced membrane potential changes in L6a FOXP2+ CT (n=16), L6b FOXP2+ CT (n=25), L6a FOXP2– CC (n=15), and L6b CC (n=12) excitatory neurons in the presence of 0.5 µM TTX to block AP firing. For statistical comparison the Wilcoxon Mann-Whitney U test was used; * p≤0.05, ** p≤0.01, *** p≤0.001, n.s., not significant. (C) Fraction of different L6 excitatory neuron types firing action potential following application of 30 µM DA in the absence of TTX. In 29% of L6b FOXP2+ CT PCs 30 µM DA induced long-lasting spike trains; this was not observed in the other L6 excitatory neuron types. (D) L6b neurons that responded with prolonged AP firing following DA (30 µM) application were located close to the L6b-WM border and showed a regular firing pattern. (E) In a subset of L6b neurons (2 out of 9, i.e. 22%), DA-induced AP firing that was independent of glutamatergic and GABAergic synaptic activity. Note the delayed onset of of AP generation. (F) The D_1_-like DA receptor antagonist SKF (1 µM) elicited a depolarisation (bottom left, see also the bar graph on the right) in all L6 excitatory neurons types showing a DA-induced depolarisation (top left, n=13).

Application of dopamine (30 µM) in the presence of TTX to L6 CT FOXP2+ excitatory neurons resulted in a depolarising response, except for two L6 CT FOXP2+ PCs which showed a weak hyperpolarisation (Membrane potential change ΔV_m_: 3.4 ± 2.4 mV, n=17 for L6a CT FOXP2+ PCs; 4.7 ± 3.2 mV, n=28 for L6b CT FOXP2+ PCs). The effect of dopamine in FOXP2– excitatory neurons in layer 6 was heterogeneous (**Figure 7B, C**): In 11 out of 16 L6a CC excitatory neurons dopamine caused a depolarisation (ΔV_m_: 2.1 ± 1.2 mV) and in the remainder a weak hyperpolarisation (ΔV_m_: −1.4 ± 1.2 mV, n=5). Most L6b CC (FOXP2–) PCs (8 out of 13) dopamine induced a hyperpolarisation (ΔV_m_: −1.7 ± 0.8 mV) while five showed a weak depolarising response (ΔV_m_: 1.0 ± 0.5 mV).

When recorded in the absence of TTX, 30 µM dopamine induced a depolarisation followed by prolonged high-frequency AP firing in 29% of L6b FOXP2+ PCs (see **Figure 7A,C**). These neurons exhibited a dendritic and axonal projection pattern similar to CT PCs, with apical dendrites terminating in L5 (**Figure 7D**) (Marx and Feldmeyer, 2013). They were located near the WM, which is significantly deeper than L6b neurons with subthreshold DA responses (relative depth in layer 6: 0.97±0.20 and 0.92±0.50, respectively) (**Figure 7D**). In all other L6 excitatory neurons, dopamine did not induce AP firing (**Figure 7A,C**). To investigate whether AP firing resulted from a dopamine-induced enhancement of synaptic input, dopamine was applied while blocking excitatory and inhibitory synaptic transmission with 10 µM CNQX, 50 µM AP5, and 10 µM gabazine. Under this condition, dopamine caused only a weak depolarisation in most L6b CT PCs; however, in two out of nine L6b CT PCs (*i.e.* 22%), AP firing was still observed (**Figure 7E**) suggesting that these neurons are strongly responding to dopamine via an intrinsic, postsynaptic mechanism independent of synaptic input. Only in one case, dopamine-induced AP firing was blocked by application of glutamate and GABA_A_ receptor antagonists.

To determine the receptor subtype mediating the dopamine-induced depolarisation, we applied the D_1_-like dopamine receptor agonist SKF81297. L6a and L6b excitatory neurons which showed a dopamine-induced depolarisation, were also depolarised by a subsequent application of 1 µM SKF81297 (**Figure 7F**). In contrast, when dopamine application resulted in a hyperpolarisation of a L6 excitatory neuron, SKF81297 showed no effect. When dopamine was applied together with 10 µM SCH23390, a D_1_ receptor antagonist, a dopamine-induced depolarisation was no longer observed suggesting that in layer 6, all FOXP2+ PCs and a subset of L6a and L6b CC neurons express D_1_-like receptors.

## Discussion

Here, we demonstrate distinct structural and functional properties of FOXP2+ and FOXP2– excitatory neurons in layer 6a and 6b of the primary somatosensory cortex. FOXP2 immunolabelling was found only in L6a and L6b CT PCs whereas all other excitatory neurons in layer 6a and 6b were FOXP2–, including L6a CC PCs, tall, putative claustrum-innervating L6a PCs, or the diverse excitatory neuron types in layer 6b (Cotel et al., 2018; Egger et al., 2020; Marx and Feldmeyer, 2013; Zhang and Deschênes, 1997). FOXP2+ CT PCs and FOXP2– excitatory neurons were found to have different synaptic dynamics and were differentially modulated by ACh and dopamine. Notably, both neurotransmitters caused prolonged AP firing in L6b FOXP2+ CT PCs, underscoring their critical role in modulating corticothalamic feedback.

### CT PC types in neocortical layer 6

Our data suggest the existence of at least three distinct subtypes of CT PCs in layer 6 of the somatosensory barrel cortex. These subtypes all exhibit FOXP2 immunoreactivity but display differential innervation of the somatosensory thalamic nuclei. The first-order VPM nucleus relays sensory signals to the neocortex via the direct, ‘lemniscal’ pathway and is only innervated by CT PCs located mostly in superficial layer 6a (Bourassa and Deschênes, 1995; Whilden et al., 2021; Zhang and Deschênes, 1997). The higher-order POm complex is part of the indirect ‘paralemniscal’ pathway and receives inputs from three distinct PC types, namely deep L5 thick-tufted PCs (Groh et al., 2008; for a review see Mease and Gonzalez, 2021; Veinante et al., 2000), a subset of PCs in deep layer 6a that innervates both VPM and POm (Bourassa et al., 1995; Whilden et al., 2021), and L6b PCs (Bourassa et al., 1995; Hoerder-Suabedissen et al., 2018).

Deep L5 CT PCs form giant synapses with POm neurons where a single presynaptic AP can trigger a burst of postsynaptic APs, effectively ‘driving’ POm neurons (Adamantidis et al., 2007; Groh et al., 2008; Reichova and Sherman, 2004). These neurons also exhibit marked short-term synaptic depression. Extracortical synaptic connections between the brainstem and thalamus show similar synaptic dynamics and are also considered as thalamic ‘drivers’ (Mo et al., 2017; Ward, 2013).

In contrast, CT PCs in cortical layer 6 form small, ‘en passant’ synaptic contacts with VPM and/or POm neurons. These CT synapses exhibit short-term synaptic facilitation and are of low efficacy (Bourassa and Deschênes, 1995; Bourassa et al., 1995; Reichova and Sherman, 2004). Therefore, CT PCs in layer 6 are considered to be ‘use-dependent’ modulators of thalamic activity. It has been suggested that the underlying mechanism is the differential synaptic release probability at different CT synapses. This probability is high for ‘drivers’ (i.e. deep L5 input to POm) but low for L6a and L6b inputs (Bickford, 2015; Sherman and Guillery, 2002; Usrey and Sherman, 2019). Previous studies have demonstrated that in layer 6, presynaptic CT PCs determine the short-term plasticity and release probability of all their synapses, whether they are intracortical or corticothalamic (Beierlein and Connors, 2002; Kim et al., 2014). Therefore, it has been argued that synaptic dynamics of PCs generally reflect their long-range axonal projection targets (Blackman et al., 2013; Brown and Hestrin, 2009). Consistent with this, we have shown that intracortical synaptic connections of L6a CT PCs but not of CC PCs or corticoclaustral PCs exhibit pronounced short-term facilitation (Yang et al., 2022). L6b CT PC synaptic connections with other neurons exhibit low synaptic efficacy, short-term synaptic facilitation, and a low release probability (Golshani et al., 2001; Lee and Sherman, 2008; Nersisyan et al., 2021; Zolnik et al., 2023; this study). This supports the notion that, like L6a CT PCs, L6b CT PCs modulate the activity of thalamic neurons.

### FOXP2 is a highly specific marker of CT neurons in layer 6a and 6b

Foxp2 is expressed in layers 6a and 6b across all cortical areas. L6a FOXP2+ neurons in rodent primary somatosensory (S1), visual (V1), and auditory (A1) cortices are CT PCs, as shown by retrograde tracer injections into first-order thalamic nuclei (Clayton et al., 2021; Kast et al., 2019; Sundberg et al., 2017; Tasic et al., 2016). Single-cell reconstructions in rat S1 barrel cortex revealed that upper L6a PCs project exclusively to the VPM, while deep L6a PCs innervate both VPM and POm, the first and higher-order thalamic nucleus (Bourassa and Deschênes, 1995; Bourassa et al., 1995; Killackey and Sherman, 2003; Whilden et al., 2021; Zhang and Deschênes, 1997).

Our findings show that FOXP2 immunoreactivity is a selective marker for all three L6 CT PC types, regardless of their projection targets (VPM alone, both VPM and POm, or POm alone). In both layers 6a and 6b, FOXP2 specifically labels typical PCs with an apical dendrite and a narrow axonal domain, but not ‘inverted’ PCs or multipolar excitatory neurons with long-range axonal projections throughout layer 6 and deep layer 5 (Egger et al., 2020; Marx and Feldmeyer, 2013; Zhang and Deschênes, 1997) or corticoclaustral PCs (Cotel et al., 2018; van Aerde and Feldmeyer, 2015; Yang et al., 2022).

The neurotensin receptor 1 gene, *Ntsr1*, is highly expressed in L6a CT PCs that project to either VPM alone or both VPM and POm (Harris et al., 2019; Whilden et al., 2021). These two L6a CT PC types have been identified in earlier retrograde tracing studies (Bourassa and Deschênes, 1995; Zhang and Deschênes, 1997). Transgenic *Ntsr1*-Cre mouse lines are utilised to identify L6a CT PCs in studies cortico-thalamo-cortical signalling (Bortone et al., 2014; Frandolig et al., 2019; Gong et al., 2007; Ibrahim et al., 2021; Kirchgessner et al., 2020; Pauzin and Krieger, 2018; Williamson and Polley, 2019). However, *Ntsr1* is also expressed in white matter neurons located directly below layer 6b (Sundberg and Granseth, 2018) and in nucleus accumbens-projecting L2/3 and L5a PCs (Babiczky and Matyas, 2022). NTSR1-mediated depolarisations have been recorded in L6b excitatory neurons and fast-spiking interneurons (Wenger Combremont et al., 2016a, b), indicating that *Ntsr1* expression is not specific to L6a CT PCs (Wenger Combremont et al., 2016a, b). Our data show that FOXP2 does not label CC excitatory neurons, consistent with a previous study (Kast et al., 2019).

In non-sensory cortical areas, *Syt6* (synaptotagmin 6 gene) is a more reliable CT neuron marker than *Ntsr1* (Vaasjo et al., 2022). *Syt6* and *Ntsr1* show minimal overlap, suggesting they label specific subsets of L6a CT PCs. *Foxp2*, however, is expressed in layer 6 of all cortical areas, indicating its ubiquity as a marker for L6 CT neurons.

In layer 6b, a distinct population of CT PCs expressing *Drd1* (dopamine receptor 1 gene) projects exclusively to the higher-order POm nucleus (Hoerder-Suabedissen et al., 2018) and is highly sensitive to the wake-promoting neuropeptide orexin (Hay et al., 2015; Zolnik et al., 2023). We identified L6b FOXP2+ PCs that innervate exclusively POm and show a robust dopamine D_1_ receptor response, indicating co-expression of *Foxp2* and *Drd1*. Nonetheless, D_1_ receptor-mediated depolarisations are also observed in L6a CT PCs and a number of L6 CC excitatory neurons suggesting that the expression of *Drd1* is not exclusive to L6b CT PCs.

### Cholinergic neuromodulation of FOXP2+ and FOXP2– excitatory neurons in layer 6a and 6b

Our findings highlight ACh as a potent modulator of both L6a and L6b FOXP2+ CT PCs, acting through muscarinic and nicotinic receptors to induce strong depolarisations even at micromolar concentrations. This modulation, which is critical during wakefulness, attention, and REM sleep, as ACh released from the basal forebrain and brainstem cholinergic nuclei can induce prolonged AP firing in almost two-thirds of L6a CT PCs and more than 75% of L6b CT PCs. In contrast, ACh consistently causes hyperpolarisation in L6a FOXP2– CC PCs, whereas L6b FOXP2– CC excitatory neurons show moderately strong ACh-induced depolarisations. This differential response suffuses a cell-specific cholinergic regulation of L6 neuron activity.

In addition, a recent study identified excitatory subplate neurons termed ‘ACh super-responders’ due to their pronounced ACh-induced depolarisations (Venkatesan et al., 2022), a feature resembling the L6b CT PCs described here. Layer 6b is considered to be a remnant of the subplate (Hoerder-Suabedissen and Molnar, 2012; Marx et al., 2017; Reep, 2000; Torres-Reveron and Friedlander, 2007), which suggests that L6b excitatory neurons retain the high ACh responsiveness akin to the ‘ACh super-responder’ subplate neurons during early postnatal development.

In situ hybridisation and antibody labelling studies have demonstrated that in layer 6, CT PCs predominantly express (α_4_)_2_(β_2_)_2_α_5_ nAChRs (Heath et al., 2010), a finding supported by several functional studies (Hay et al., 2016; Kassam et al., 2008; Koukouli et al., 2017; Venkatesan et al., 2023; Yang et al., 2024). This α_4_β_2_α_5_ nAChR is characterised by high ACh sensitivity and slow desensitisation, resulting in a large and sustained ACh response in these neurons (Kuryatov et al., 2008; Ramirez-Latorre et al., 1996). In addition, α_4_β_2_α_5_ nAChRs exhibit a high Ca^2+^ permeability, which facilitates modulation of presynaptic neurotransmitter release and significantly increases synaptic efficacy. As previously demonstrated, ACh enhances synaptic release at L6a CT PC synapses, with the activation of presynaptic nAChRs being identified as a key mechanism in this process (Yang et al., 2020). The findings presented here indicate a comparable effect at the L6b CT pyramidal cell synapse.

Furthermore, the activation of postsynaptic nAChRs also significantly enhances the probability of AP firing in L6a and L6b CT PCs. The cholinergic modulation of L6 CT neurons demonstrated here implies that even relatively low ACh concentrations can efficiently recruit VPM and POm neurons, effectively modulating the gain of thalamocortical output. The prolonged firing can enhance thalamocortical feedback, crucial for sensory signal amplification during states of heightened attention (Hasselmo and Sarter, 2011).In particular, the strong ACh response in L6b FOXP2+ CT PCs suggests that they play a role in modulating thalamic activity dynamically, which could be essential for maintaining sensory signal fidelity during behavioural states requiring high attentiveness.

### Dopaminergic neuromodulation of FOXP2+ and FOXP2– excitatory neurons in layer 6a and 6b

Dopaminergic afferents to the neocortex originate mainly from the ventral tegmental area (VTA) and, to a lesser extent, the substantia nigra pars compacta (Björklund and Dunnett, 2007; Descarries et al., 1987). The somatosensory cortex receives dopaminergic inputs from mesocortically and mesocorticolimbically projecting VTA neurons (Aransay et al., 2015). In rats, L6 CT PCs express dopamine D_1_ receptors as indicated by the presence of the dopamine- and cAMP-regulated neuronal phosphoprotein (DARPP-32; Ouimet, 1991). Dopamine D_1_ receptor-expressing neurons are primarily located in deep cortical layers 5 and 6 (Anastasiades et al., 2019; Dembrow and Johnston, 2014; Gaspar et al., 1995; Santana and Artigas, 2017), where the density of dopaminergic afferents is highest in rodents (Berger et al., 1991). However, in PFC, the majority of *Drd1*-expressing neurons were L6 CC neurons, including inverted PCs (Anastasiades et al., 2019). In contrast, in the somatosensory cortex, the largest dopamine D_1_ receptor response was observed in L6b CT PCs, resulting AP firing in nearly 30% of the tested neurons. A subset of L6 FOXP2– CC excitatory neurons also exhibited a weak depolarising response to dopamine, which was smaller than in L6 FOXP2+ CT PCs. This indicates that *Drd1* expression is probably not exclusive to L6b CT PCs projecting to the POm. Nevertheless, the high expression of *Drd1* and the density of functional dopamine D_1_ receptors in L6b CT PCs may be the reason that this gene is considered to be a specific marker for these neurons.

The robust dopamine responses observed in L6b FOXP2+ CT PCs are consistent with the known roles of dopamine in modulating cortical activity related to cognition and attention (Nieoullon, 2002; Robbins and Arnsten, 2009; Speranza et al., 2021). Our findings that dopamine induces prolonged AP firing in L6b FOXP2+ CT PCs suggest that these neurons could play a significant role in attention regulation and sensory information processing, particularly through the POm. The capacity of dopamine to enhance the excitability of these neurons could facilitate a state-dependent modulation of sensory processing, thereby reinforcing the link between dopaminergic signalling and cognitive functions such as attention and working memory.

### Role of L6a and L6b CT PCs in corticothalamic signalling

In the somatosensory system, L6a CT PCs innervate not only excitatory neurons in the VPM (upper layer 6a) or VPM and POm (lower layer 6a) but also inhibitory neurons in the thalamic reticular nucleus (TRN) which projects directly to both VPM and POm (Pinault, 2004). As a result, L6a CT PCs modulate the activity of these thalamic nuclei by inducing direct, monosynaptic excitation and subsequent disynaptic inhibition through TRN neurons (Halassa and Acsady, 2016; Lam and Sherman, 2010). By contrast, L6b CT PCs exclusively provide direct facilitating synaptic input to the POm without recruiting the TRN neurons, and thus do not engage this intrathalamic feedforward inhibitory circuit (Bourassa et al., 1995; Crandall et al., 2015; Hoerder-Suabedissen et al., 2018). This L6b POm input is powerfully excited by orexin, a neuropeptide crucial for wakefulness and attention, which drives wake-like cortical states (Bayer et al., 2004; Hay et al., 2015; Zolnik et al., 2023). Orexin depolarises L6b CT PCs that are also particularly sensitive to ACh and dopamine, two neurotransmitters that are also involved in wakefulness and attention, as shown here. In concert, these neuromodulators will mediate a marked activation of the POm.

ACh and dopamine also trigger the activation of L6a CT PCs, enhancing VPM activity. However, ACh suppresses TRN activity through mAChRs (Lam and Sherman, 2010), reducing net inhibition by L6 CT circuits (Olsen et al., 2012). Additionally, ACh-induced L6b CT neuron activation recruits large L6 translaminar, fast-spiking interneurons whose axons span almost the entire cortical depth and primarily impact L2/3 and L5 PCs (Bortone et al., 2014; Zolnik et al., 2023). Therefore, predicting the overall effect of ACh-induced firing in L6a CT PCs is challenging and requires further study because ACh enhances the excitability of several neuron types in layer 6 (Obermayer et al., 2017; Radnikow and Feldmeyer, 2018).

Previous research demonstrated that posterior parietal cortex (POm) stimulation enhances and prolongs cortical sensory signals, likely due to increased thalamic activity leading to elevated thalamic input to the neocortex (Mease et al., 2016). Our data indicate that ACh may regulate this process, thereby increasing the gain of POm output. This could serve to amplify relevant sensory input during heightened attention and arousal.

Dopamine release plays a crucial role not only in motor functions but also in cognition and attention (Nieoullon, 2002). Since dopamine specifically enhances the activity of the L6b-POm pathway, it is likely to increase POm activity similarly to acetylcholine. However, the practical implications of dopaminergic input for cortico-thalamocortical signalling are not yet fully understood and require further study.

### Conclusion

This study identifies three distinct FOXP2+ CT PC types in layer 6 that are tightly regulated by neuromodulators. Our findings reveal distinctive properties of these neurons, particularly L6b CT PCs, which are highly responsive to neuromodulators. The results highlight the important role of L6b CT PCs in regulating the cortico-thalamo-cortical feedback loop. By identifying specific subtypes of FOXP2+ CT PCs and their unique synaptic and neuromodulatory properties, this study advances our understanding of the neural mechanisms involved in sensory processing, attention and arousal. It may also inform future research into therapeutic targets for neurological disorders involving corticothalamic dysfunction.

The distinct roles of L6a and L6b CT PCs in modulating thalamic activity through direct and indirect pathways highlight their importance in sensory processing under different behavioural states. L6a CT PCs modulate both excitatory and inhibitory thalamic circuits, potentially affecting sensory gating and attentional mechanisms. In contrast, L6b CT PCs provide direct excitatory input to the POm, which is modulated by ACh and dopamine. This input is critical for enhancing sensory input during wakefulness and attention. This division of labour within L6 CT PCs underscores the complexity of corticothalamic interactions and their relevance to adaptive sensory processing.

## Materials and Methods

All experimental procedures involving animals were performed in accordance with the guidelines of the Federation of European Laboratory Animal Science Association (FELASA), the EU Directive 2010/63/EU, and the German animal welfare law. Official licence for ‘in vivo’ experiments were granted by the ‘Landesamt für Natur- und Verbraucherschutz’ of the federal state of Northrhine-Westphalia.

### Slice preparation

In this study, Wistar rats (Charles River, either sex) aged 16–24 postnatal days (PDs) were anaesthetised with isoflurane at a concentration < 0.1% and decapitated. The brain was quickly removed and placed in an ice-cold modified artificial cerebrospinal fluid (ACSF) containing a high Mg^2+^ and a low Ca^2+^ concentration (4 mM MgCl_2_ and 1 mM CaCl_2_) to reduce potentially excitotoxic synaptic transmission during slicing; other components were identical to those in the perfusion ACSF as described below. To maintain adequate oxygenation and a physiological pH level, the solution was constantly bubbled with carbogen gas (95% O_2_ and 5% CO_2_). Thalamocortical slices (Agmon and Connors, 1991; Feldmeyer et al., 1999) were cut at 350 µm thickness using Leica VT1000S vibrating blade microtome and then transferred to an incubation chamber containing preparation solution for a recovery period of 30-60 minutes at room temperature before being transferred to the recording chamber.

### Solution

During whole-cell patch-clamp recordings, slices were continuously superfused (perfusion speed ∼5 ml/min) with ACSF containing (in mM): 125 NaCl, 2.5 KCl, 1.25 NaH_2_PO_4_, 1 MgCl_2_, 2 CaCl_2_, 25 NaHCO_3_, 25 D-glucose, 3 mho-inositol, 2 sodium pyruvate and 0.4 ascorbic acid, bubbled with carbogen gas and maintained at 30-33 °C. Patch pipettes were pulled from thick-wall borosilicate glass capillaries and filled with an internal solution containing (in mM): 135 K-gluconate, 4 KCl, 10 HEPES, 10 phosphocreatine, 4 Mg-ATP, and 0.3 GTP (pH 7.4 with KOH, 290-300 mOsm). The ‘searching’ pipette was filled with an internal solution in which K^+^ is replaced by Na^+^ (containing (in mM): 105 Na-gluconate, 30 NaCl, 10 HEPES, 10 phosphocreatine, 4 Mg-ATP and 0.3 GTP), in order to prevent the depolarisation of neurons during searching for presynaptic neurons. Biocytin at a concentration of 5 mg/ml was added to the internal solution in order to stain patched neurons after recordings. In addition, biocytin-conjugated Alexa Fluor 594 dye (1:500, Invitrogen) was added to the internal solution for *post hoc* identification of patched neurons during fluorescence imaging.

### Electrophysiological recording and analysis

The slices and neurons were visualised using an upright microscope equipped with an infrared differential interference contrast (IR-DIC) optics. The barrels in L4 of the somatosensory cortex can be identified as dark stripes with light ‘hollows’ at low magnification (4x objective) and were visible in 6-8 consecutive slices (Feldmeyer et al., 1999). For single-cell recordings, neurons throughout the entire L6, i.e. from the border with deep L5b to the WM were randomly selected (Marx and Feldmeyer, 2013; Qi and Feldmeyer, 2016). L6 pyramidal cells (PCs) and L6 interneurons can be distinguished by their soma shape an the presence or absence of an ‘upright’ apical dendrite at high magnification (40x objective). They can also be differentiated by their intrinsic action potential (AP) firing patterns and - following histological staining - by their morphology.

Whole-cell patch clamp recordings were made using an EPC10 amplifier (HEKA, Lambrecht, Germany). Signals were sampled at 10 kHz, filtered at 2.9 kHz using Patchmaster software (HEKA), and later analysed off-line using Igor Pro software (Wavemetrics, USA). Recordings were performed using patch pipettes of 5-8 MΩ resistance. To find synaptic connections we used a ‘searching procedure’ described previously (Feldmeyer et al., 1999; Qi et al., 2015). Briefly, after patching a ‘postsynaptic neuron’, potential presynaptic neurons were patched with a ‘searching’ pipette filled with a Na-based internal solution. When an AP elicited in ‘loose cell-attached’ mode resulted in an excitatory postsynaptic potential (EPSP) in the postsynaptic neuron, this presynaptic neuron was re-patched with a new pipette filled with biocytin-containing internal solution. EPSPs were recorded from a postsynaptic neuron by evoking 2-3 APs or a train of 10 APs in the presynaptic neuron with brief (5 ms) depolarising pulses at 10 Hz. The interval between the simulation trains was 10-20 s. For each synaptic connection, 40 or more sweeps were collected during recordings.

Custom-written macros in Igor Pro 6 (WaveMetrics, Lake Oswego, USA) were used to analyse the recorded electrophysiological signals. To assess passive and active action potential firing properties, a series of 1 s current pulses starting with initial hyperpolarisation, followed by depolarisation, were elicited under current clamp configuration. Neurons with a series resistance exceeding 40 MΩ or that displayed a depolarised membrane potential (> −55 mV) after the cell membrane was ruptured were excluded from the data analysis. The resting membrane potential (V_rest_) was recorded immediately after establishing the whole-cell recording configuration. Passive membrane properties, including the input resistance (R_in_), membrane time constant τ_m_, and voltage sag, were measured by analysing membrane potential (V_m_) traces induced by a series of hyper- and depolarising subthreshold current pulses. Additionally, for the 1^st^ AP elicited by a rheobase current step, the threshold, amplitude, half-width, latency, and AHP amplitude and latency were determined as single action potential properties.

Properties related to repetitive firing, such as spike frequency adaptation, standard deviation (s.d.) of inter-spike intervals (ISIs), AP amplitude and half-width accommodation, and AHP change, were measured for a current step that elicited approximately 10 APs. The analysis of most electrophysiological parameters has been described previously (Emmenegger et al., 2018) with two exceptions. *AHP latency* was defined as the time interval between the AP threshold and AHP trough and the *adaption ratio* was calculated by measuring the last inter-stimulus interval (ISI) relative to the third ISI (excluding ISIs of initial bursts, doublets, or triplets); *the s.d. of ISIs* was calculated as the standard deviation of ISI_3_, ISI_4_, …, ISI_9_; the AHP change was calculated as the voltage difference between the last AHP and the 1st AHP (Beierlein et al., 2003; Crandall et al., 2017). Synaptic properties were evaluated as described previously (Feldmeyer et al., 1999; Feldmeyer et al., 2002). All unitary EPSP (uEPSP) recordings were aligned to their corresponding presynaptic AP peaks, and the mean uEPSP was generated by averaging the recordings. The amplitude of the EPSP was calculated by subtracting the mean baseline from the maximum voltage of the postsynaptic event. The paired-pulse ratio was defined as the amplitude of the 2nd uEPSP divided by that of the 1st uEPSP, which was elicited by presynaptic action potentials at a stimulation frequency of 10 Hz.

### Drug application and analysis

Acetylcholine (ACh) at either low (30 µM) or high (1 mM) concentration and dopamine (DA, 30 µM) were applied via the perfusion system. Atropine (ATRO, 200 nM), mecamylamine (MEC, 10 µM), dihydro-ß-erythroidine (DHßE, 10 µM), SKF81297 (1 µM) and tetrodotoxin (TTX, 0.5 µM) were all bath applied; drugs were purchased from Sigma-Aldrich or Tocris. For single-cell recordings, a stable baseline with fluctuation in Vm <1 mV was recorded for 3 min before drug application via the perfusion system. ΔV_m_ was calculated as the difference between the peak V_m_ excursion (positive or negative) following drug application and the baseline potential. For paired recordings, 10-20 sweeps were recorded as baseline before drug application. Sweeps were continuously recorded during drug application (40-60 sweeps) and washout (80-100 sweeps).

### Immunohistochemical staining and imaging

To determine the FOXP2 immuoreactivity in L6 neurons recorded in acute brain slices, slices (350 µm) were fixed after electrophysiological recordings with 4% paraformaldehyde (PFA) in 100 mM phosphate buffered saline (PBS) for at least 24 h at 4°C and then permeabilised in 1% milk power solution containing 0.5% Triton X-100 and 100 mM PBS. Primary and secondary antibodies were diluted in the permeabilisation solution (0.5% Triton X-100 and 100 mM PBS) shortly before experiments. For FOXP2 staining of patched neurons, slices were incubated overnight with Goat-anti-FOXP2 primary antibody (1:500, Santa Cruz Biotechnology) at 4°C and then rinsed thoroughly with 100 mM PBS. Subsequently, slices were treated with Alexa Fluor secondary antibodies (1:500) for 2–3 h at room temperature in the dark. After being rinsed in 100 mM PBS, slices were embedded in Fluoromount. Fluorescence images were taken using either the Olympus CellSens platform or a Zeiss Scope A1 microscope. Prior to antibody labelling, the position of the patched neurons was marked by addition of the biocytin-conjugated Alexa 594 dye to the intracellular solution (Figure 1C,D). Slices were then incubated in 100 mM PBS overnight and processed for subsequent histological staining for morphological reconstruction and analysis as described below.

For FOXP2 antibody labelling of whole-slices, thinner slices (150 µm) were prepared and processed following the same procedure as described above. To determine the percentage of FOXP2– immunoreactive neurons in cortical L6 and the POm, slices were treated with both antibodies for FOXP2 and the neuronal marker protein NeuN. The primary antibody used for NeuN staining was mouse-anti-NeuN (1:250, Merck-Millipore). Fluorescence images (single images and image stacks) from different cortical and subcortical brain regions were acquired at constant exposure times and at different magnification levels. To ensure comparability between the co-localisation of our target antigens FOXP2 and NeuN, we maintained the same settings, slice, region, and intra-stack image distance. Single images and image stacks were assessed using the freely available ImageJ software (National Institute of Health; http://rsbweb.nih.gov). Image stacks were first visualised by a 3D surface plot and further processed by a standard z-projection. FOXP2+ and NeuN+ neurons were counted with the aid of the cell counter plugin tool for ImageJ. Counts were averaged across all slices and analysed to determine the fraction of cells in which FOXP2 and NeuN were co-localised.

To recover the morphology of biocytin-filled neurons or neuron pairs, slices were rinsed several times in 100 mM PBS and then treated with 1% H_2_O_2_ in PBS for about 20 min in order to reduce any endogenous peroxidase activity. Slices were rinsed repeatedly with PBS and then incubated in 1% avidin-biotinylated horseradish peroxidase (Vector ABC staining kit, Vector Lab. Inc., Burlingame, USA) containing 0.1% Triton X-100 for 1 h at room temperature. The reaction was catalysed using 0.5 mg/ml 3,3-diaminobenzidine (DAB; Sigma-Aldrich, St.Louis, Mo, USA) as a chromogen. Slices were then rinsed with 100 mM PBS, followed by slow dehydration with ethanol in increasing concentrations and finally in xylene for 2–4 h. After that, slices were embedded using Eukitt medium (Otto Kindler GmbH, Freiburg, Germany).

### Morphological reconstruction and analysis

Computer-assisted morphological 3D reconstructions of L6 neurons were made using NEUROLUCIDA^®^ software (MicroBrightField, Williston, VT, USA) and Olympus BV61 microscopy fitted with a 100x objective. Neurons were selected for reconstruction based on the quality of biocytin labelling when background staining was minimal. The cell body, dendritic and axonal branches were reconstructed manually under constant visual inspection to detect thin and small collaterals. Barrel and layer borders, pial surface and the white matter (WM) were delineated during reconstructions at lower magnification 4x. The position of soma and layers were confirmed by superimposing the DIC images taken during the recording. The tissue shrinkage was corrected using correction factors of 1.1 in the x–y direction and 2.1 in the z direction (*Marx et al., 2012*). Because the cortical thickness of slices where neurons were stained vary from slice to slice, individual neuronal reconstructions were normalised to the average thickness along the pia-WM axis before the morphological analysis. Parameters characterising general morphological properties of reconstructed neurons, e.g. the perimeter and area of somata, the number and length of axonal and dendritic branches, were extracted using the NEUROEXPLORER® software (MicroBrightField Inc., Willston, VT, USA). Other morphological parameters have been described before (Yang et al. 2020) except for the absolute and relative depth of somata: soma depth was calculated as the pia-to-soma distance along the pia-WM axis and the relative depth was calculate as the ratio of the pia-to-soma distance to the pia-to-WM distance.

Furthermore, the 3D density maps of axonal and dendritic length were obtained using computerised 3D reconstructions, where the length of the axonal and dendritic tree per unit volume of 50 × 50 × 50 µm^3^ was calculated. The soma centre of each neuron was given the co-ordinates of *X*, *Y*, *Z* = (0, 0, 0), and the relative coordinate of the beginning and endpoint of each segment in the tracing were obtained using the segment point analysis in NEUROEXPLORER^®^. Further steps were carried out in Matlab (MathWorks) using a custom-written algorithm. The 3D axonal and dendritic density maps were calculated for each representative reconstructed neuron from a single group. These were then averaged to obtain the 3D density map for this group. Individual density maps were aligned with respect to the barrel centre. The averaged density map for each group was smoothed using the 3D smooth function in Matlab with a Gaussian kernel (s.d.=50 µm). Isosurfaces at the 80-percentile were calculated for the smoothed density maps. Finally, axonal and dendritic density maps were visualised after projecting to 2D or 1D using two different colors, e.g. blue and red, respectively.

### Retrograde tracing in combination with FOXP2 immunolabelling

Male Wistar rats were provided by Charles River Laboratories. All experiments were carried out after evaluation by the local German authorities, and in accordance with the animal welfare guidelines of the Max Planck Society. Neurons in cortex were retrogradely labelled as described previously (Rojas-Piloni et al., 2017). Briefly, young (P22–P25) male Wistar rats were injected with 1 mg/ml buprenorphine SR (0.05ml subcuteneous) approximately 30 min prior to surgery, and then anaesthetised with isoflurane/O_2_ gas mixture (2% isoflurane v/v). Rats were then placed in a stereotaxic frame (Kopf Instruments 1900) and given an injection of 0.25% bupivacaine (0.10cc, s.q.) at the incision site. Then a 5 cm incision across the midline was made to expose the skull. Both bregma and lambda were marked with a surgical pen. Two small craniotomies were made using a dental drill (Osada EXL-M40) over the injection sites of the right cerebral hemisphere. Injection site coordinates were as follows (in mm): POm: 2.1 lateral from midline, 3.25 posterior to bregma and 5.2 deep from the pia; VPM: 3.25 lateral from midline, 2.9 posterior to bregma and 5.5 deep from the pia. Prior to injecting tracers into the VPM and POm, the head of the rat was levelled with a precision of 1 µm in both the medial–lateral and anterior–posterior planes using an electronic leveling device (eLeVeLeR; Sigmann Elektronics, Hüffenhardt, Germany) mounted to an adapter for the Kopf stereotax. Retrograde tracers, CTB-488 and CTB-647 (Molecular Probes; 1 mg/ml in PBS) were pressure injected (50-200 nl) under visual control into VPM and POm thalamus, respectively, using a 30cc syringe coupled to a calibrated glass injection capillary. After injection of tracers, the incision site was thoroughly cleaned with saline and sutured. Rats underwent a 5–7-day incubation period after tracer injection before transcardial perfusion with 4% PFA dissolved in 100 mM PBS. Brains were then cut into consecutive 40 µm thick coronal sections and immunolabelled with NeuN or FOXP2 as described above. The injection sites in VPM and POm were validated on low-resolution images of the thalamus (Figure 2A). Quantification of retrogradely labeled neurons in cortex was performed on high-resolution confocal images (Figure 2B-D). The images were acquired using a confocal laser scanning system (Leica Application Suite Advanced Fluorescence SP5; Leica Microsystems) with glycerol immersion objectives (HC PL APO 10x 0.4 N.A., HC PL APO 20x 0.7 N.A.), a tandem scanning system (Resonance Scanner: 8 kHz scanning speed), spectral detectors with hybrid technology (GaAsP photocathode; 8x line average) and mosaic scanning software (Matrix Screener, beta version provided by Frank Sieckmann, Leica Microsystems). The laser excitation/emission settings were: AlexaFluor-488 (excitation: 488nm (Argon-laser); emission detection range (495-550 nm), AlexaFluor-594 (excitation: 561 nm (DPSS-laser); emission detection range (600-630 nm), AlexaFluor-647 (excitation 633 nm (HeNe-laser); emission detection range (650-785 nm). For CTB-488 and CTB-647 double injection experiments, done in combination with FOXP2 staining conjugated with Alexa 594, the images were taken sequentially using either the 10x objective with a digital zoom of 1.7 (0.868 x 0.868 µm per pixel), or a 20x objective with a digital zoom of 2.0 (0.361 x 0.361 µm per pixel).

### Statistical analysis

For all data, the mean ± s.d. was given. Wilcoxon Mann-Whitney U test was performed to access significant difference between individual groups. Wilcoxon signed rank test was used to compare paired data. Statistical significance was set at P < 0.05, n indicates the number of neurons/pairs analysed. To prepare box plots, the web application PlotsOfData (https://huygens.science.uva.nl/PlotsOfData/) was used (Postma and Goedhart, 2019).

## Acknowledgement

We would like to thank Werner Hucko for excellent technical assistance. We thank Dr. Karlijn van Aerde for custom-written macros in Igor Pro software. We are grateful for funding support from the European Union’s Horizon 2020 Framework Programme for Research and Innovation under the Framework Partnership Agreement No. 650003 (HBP FPA) to D.F.

