## Supplemental Figures and Tables for "FOXP2-immunoreactive, corticothalamic pyramidal cells in neocortical layers 6a and 6b are tightly regulated by neuromodulatory systems"

### Supplemental Material

Guanxiao Qi[1*], Danqing Yang[1,2*], Felipe Yañez[3], Fernando Messore[3], Arco Bast[3], Marcel Oberlaender[3,4] and Dirk Feldmeyer[1,2,5]

Guanxiao Qi[1*], Danqing Yang[1,2*], Felipe Yañez[3], Fernando Messore[3], Arco Bast[3], Marcel Oberlaender[3,4] and Dirk Feldmeyer[1,2,5]

[1] Institute of Neuroscience and Medicine 10, Research Centre Jülich, 52425 Jülich, Germany

[2] Dept. of Psychiatry, Psychotherapy and Psychosomatic, RWTH University Hospital, 52074 Aachen, Germany

[3] In Silico Brain Sciences Group, Max-Planck-Institute for Neurobiology of Behavior – caesar, 53175 Bonn, Germany

[4] Department of Integrative Neurophysiology, Center for Neurogenomics and Cognitive Research, Vrije Universiteit Amsterdam; 1081 Amsterdam, the Netherlands

[5] Jülich-Aachen-Research Alliance 'Brain' - Translational Brain Medicine

*these authors have contributed equally to this work

**Correspondence:**

Univ.-Prof. Dr. Dirk Feldmeyer
Forschungszentrum Jülich GmbH
Institut für Neurowissenschaften und Medizin 10 (INM-10)
Leo-Brandt-Strasse
52425 Jülich
Germany


**Suppl. Figure S1**

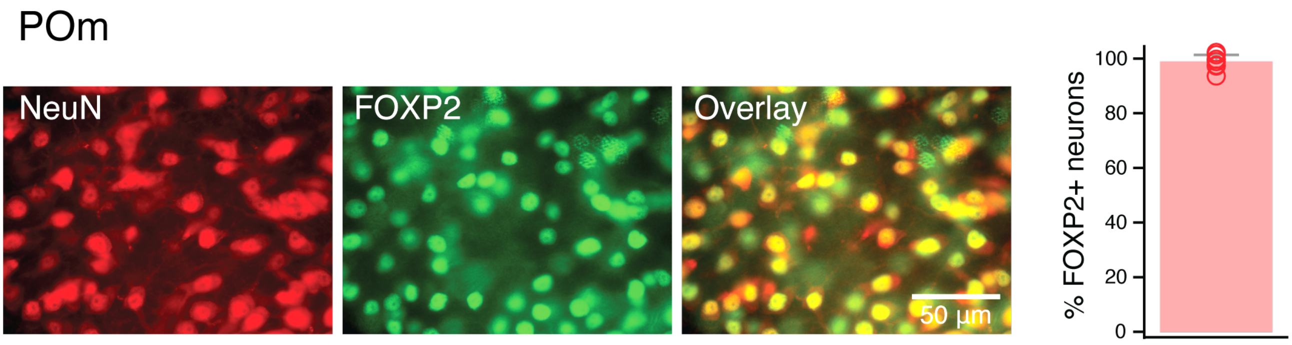

**Suppl. Figure S1**. FOXP2 Immunoreactivity in the POm. Virtually all neurons in this thalamic nucleus are FOXP2 immunoreactive.

**Suppl. Figure S2**

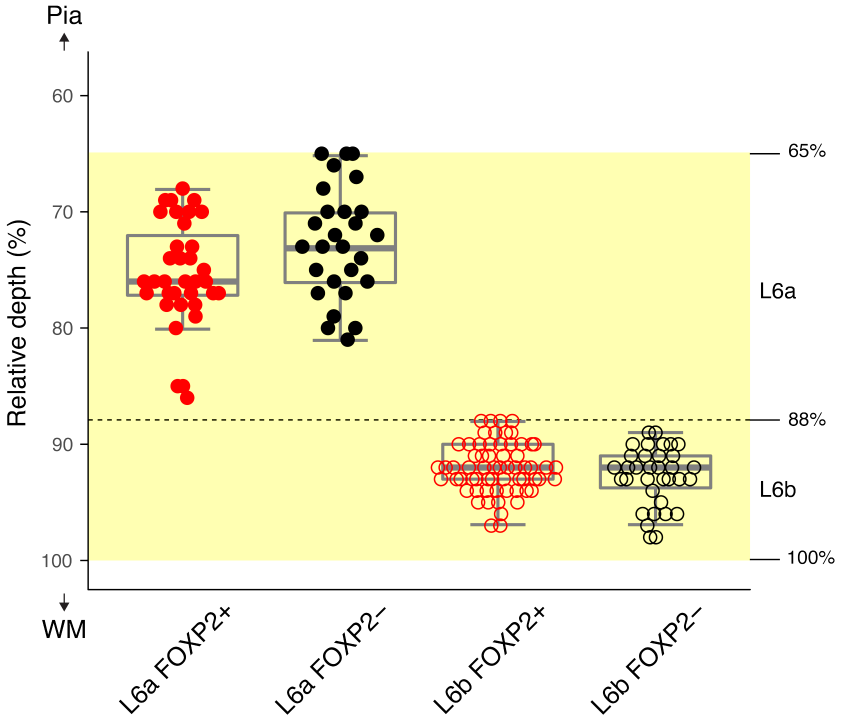

**Suppl. Figure S2**. Relative somatic depth of recorded L6a FOXP2+ (n=35), L6a FOXP2– (n=27), L6b FOXP2+ (n=63) and L6b FOXP2– (n=34) excitatory neurons. Layer 6 comprises approximately one third of the entire cortical depth (between 65% and 100% from the Pia (0%) to the layer6/WM border (100%)); it is highlighted as the semi-transparent yellow zone. Layer 6a is located in the upper 2/3 while L6b in the lower third (between 88% and 100%) of L6.

**Suppl. Figure S3**

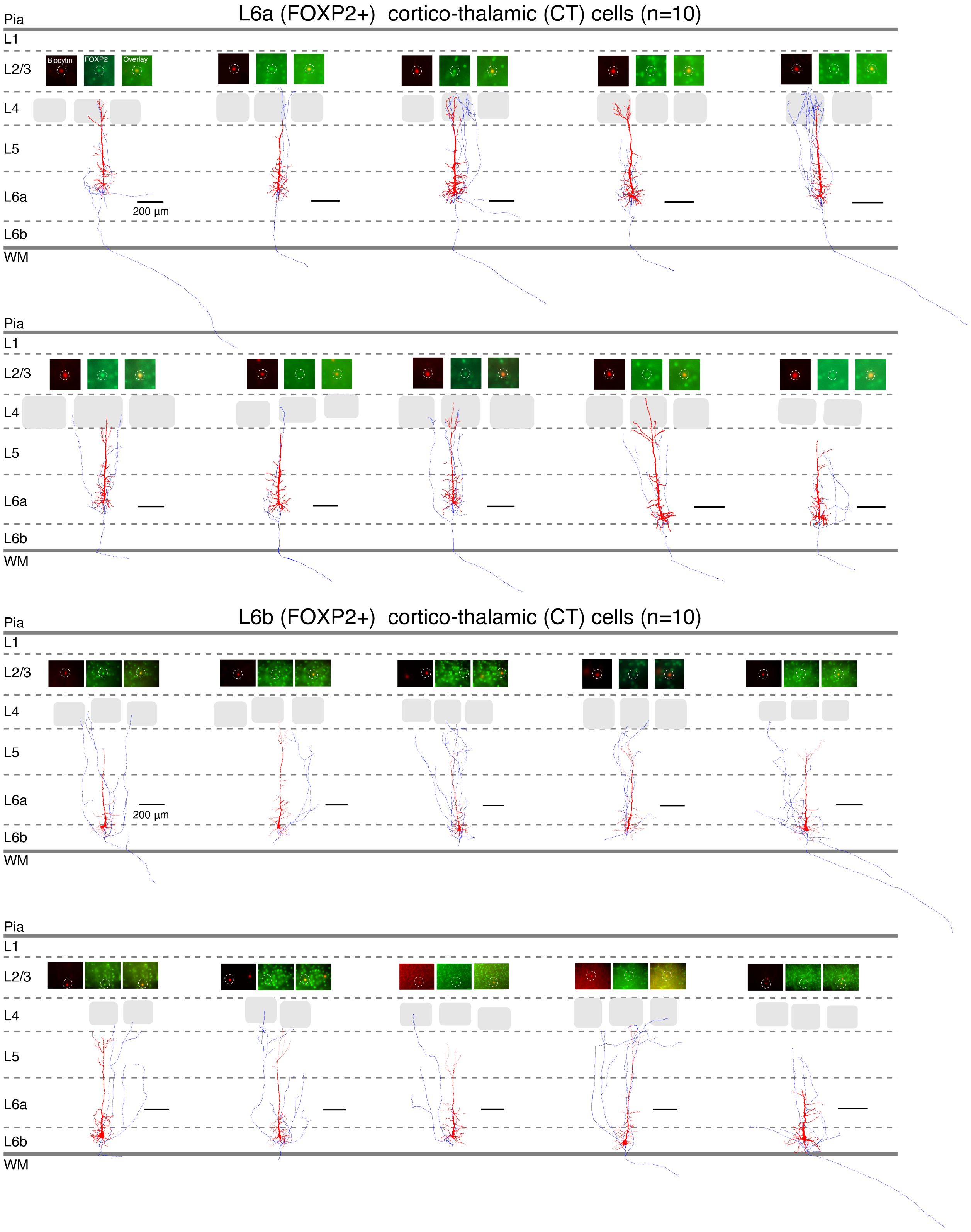

**Suppl. Figure S3**. Gallery of L6a FOXP2+ (n=10), L6b FOXP2+ (n=10), and L6a FOXP2– (n=10) and L6b FOXP2– (n=10) excitatory neuron morphologies. Insets, FOXP2 immunoreactivity.

**Suppl. Figure S3** (continued)

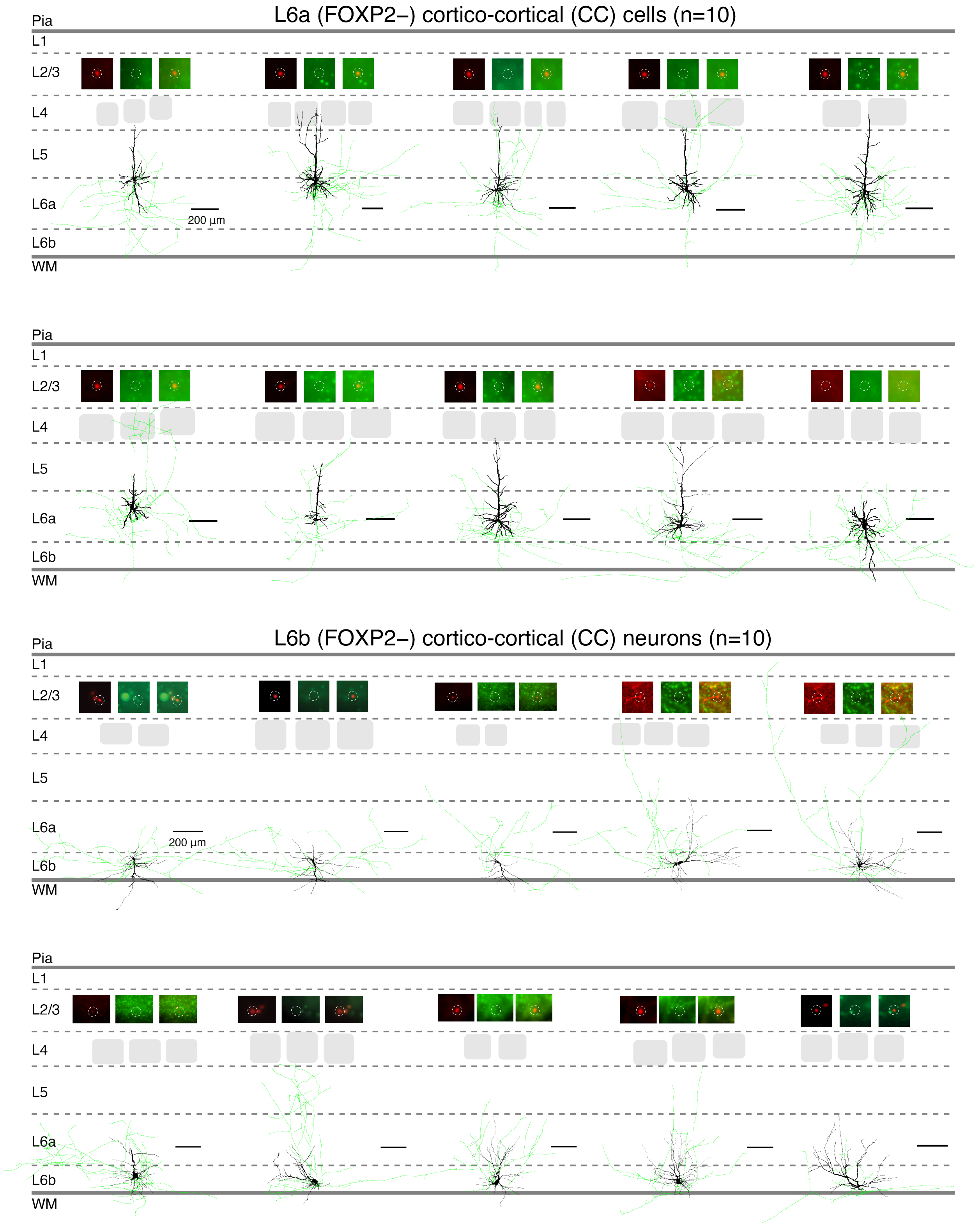

**Suppl. Figure S4**

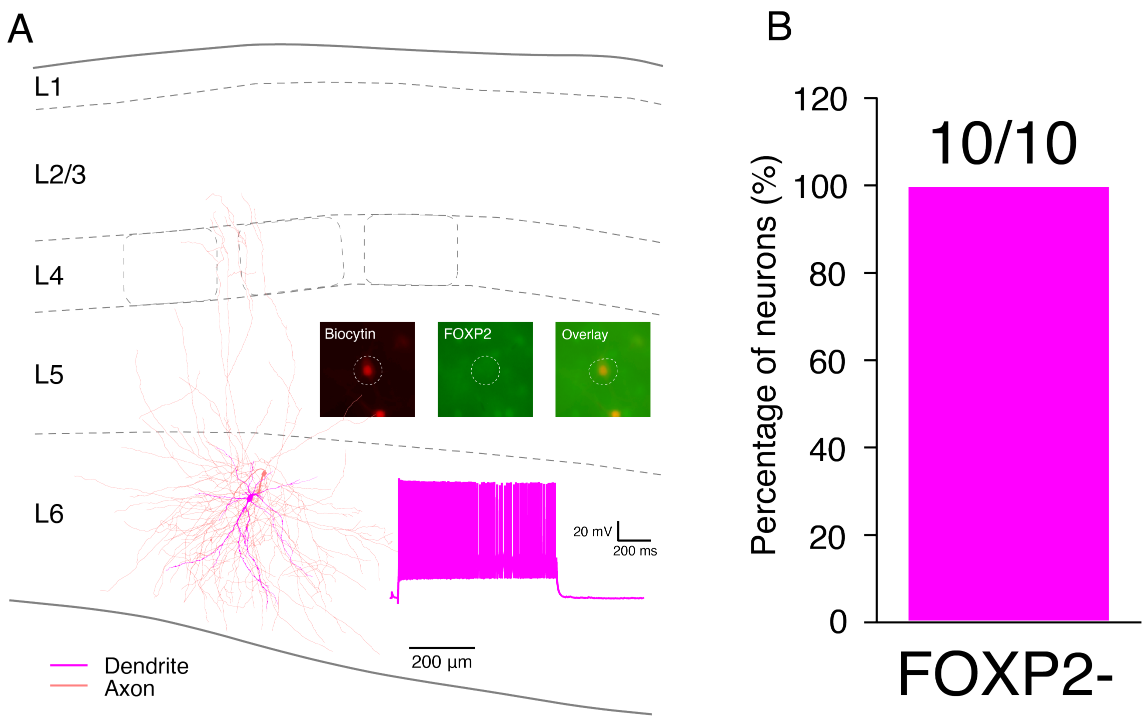

**Suppl. Figure S4**. L6 interneurons show no FOXP2 immunolabelling. (A) Morphological reconstruction of a L6 FOXP− fast spiking interneuron. (B) Ten L6 interneurons including 3 fast spiking and 7 non-fast spiking interneurons were examined for FOXP2 labelling. None of them was FOXP2 immunoreactive (see insets in A).

**Suppl. Figure S5**

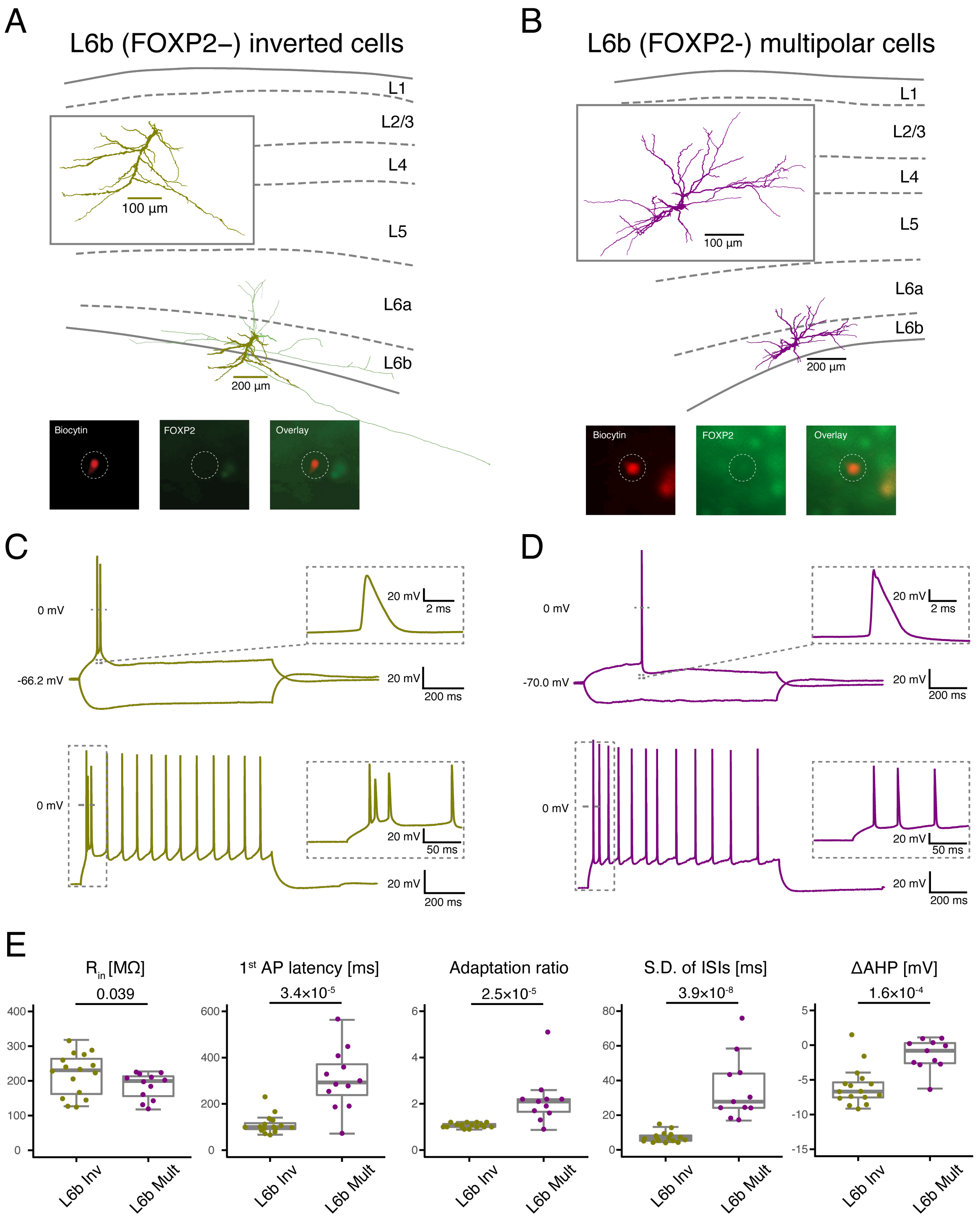

**Suppl. Figure S5**. L6b FOXP2– inverted pyramidal and multipolar spiny neurons show distinct electrophysiological properties. (A) Representative morphological reconstructions (upper) of a L6b inverted pyramidal cell and (B) a L6b multipolar spiny neuron (right), both of which are not FOXP2+ (bottom photographs). (C, D) Electrophysiological recordings of AP firing patterns from the same neurons shown in (A, B). (E) Box plots for electrophysiological properties of L6b inverted (n=17) and multipolar (n=13) spiny neurons. The box indicates the interquartile range (IQR), the whiskers show the range of values that are within 1.5*IQR and a horizontal line indicates the median. P value was calculated using the Wilcoxon Mann-Whitney U test.

**Suppl. Figure S6**

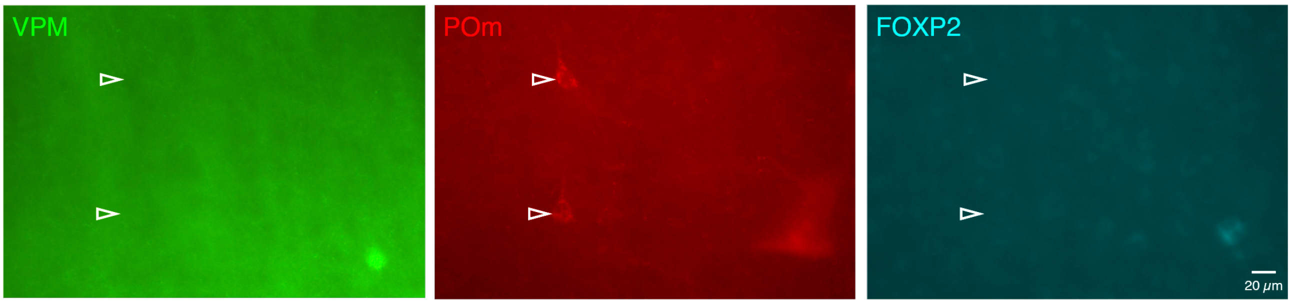

**Suppl. Figure S6**. L5b pyramidal cells were not retrogradely labelled by CTB injections in the VPM (left, green) but only in the POm (middle, red). The two L5b pyramidal neurons innervating the POm do not show FOXP2 immunoreactivity (right, blue). Individual retrogradely labelled pyramidal cells are marked by white open arrow heads.

**Suppl. Figure S7**

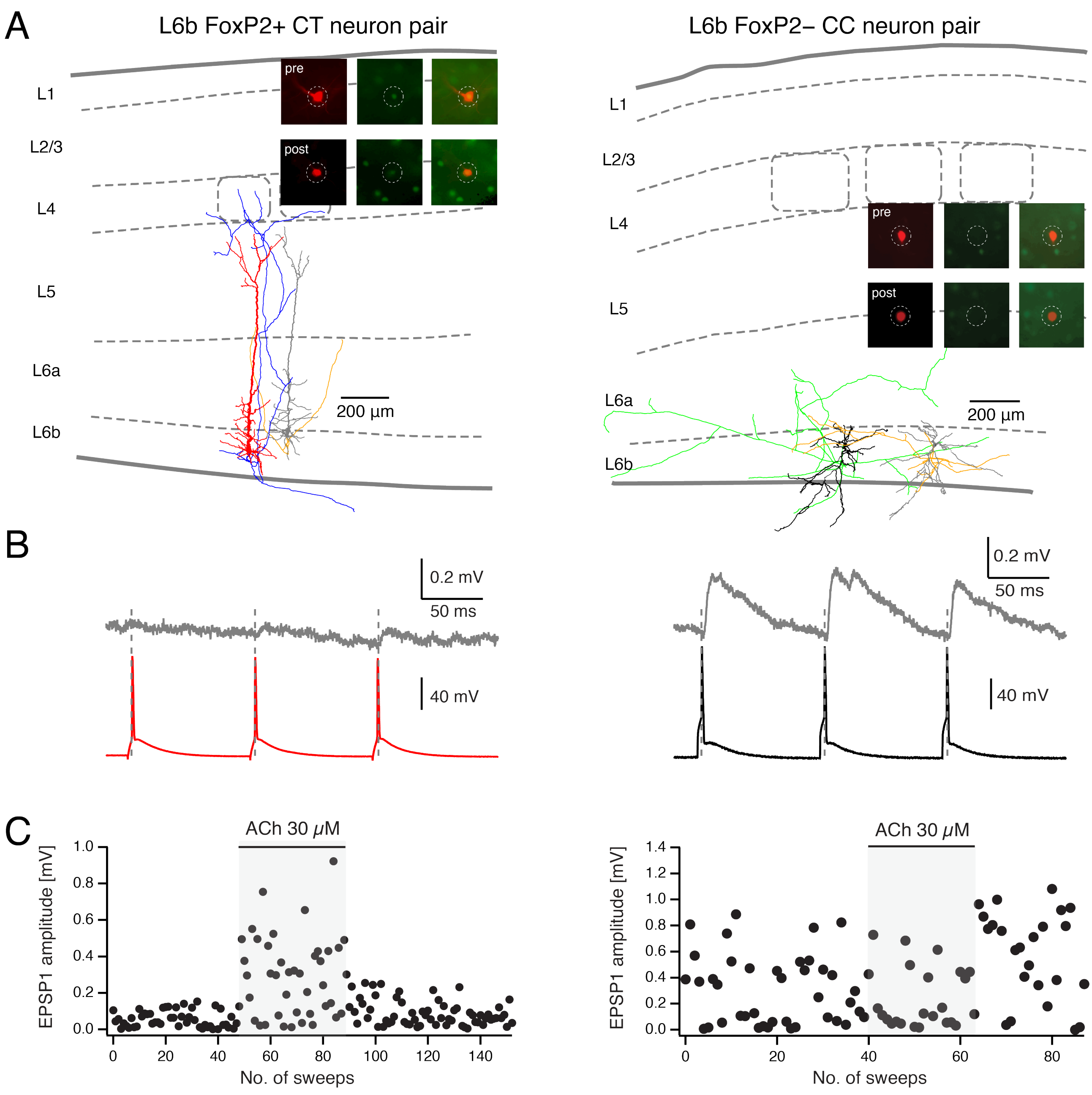

**Suppl. Figure S7**. Differential cholinergic modulation of synaptic connections established by L6b FOXP2+ and FOXP2– excitatory neurons. (A) Morphology and FOXP2 immunoreactivity (insets) for a L6b FOXP2+ (left) and FOXP2– (right) neuron pair. (B) EPSPs (upper) recorded under control condition in postsynaptic neurons induced by three APs (lower) in presynaptic neurons from the same neuron pairs shown in (A). (C) Time course of 1st EPSP amplitude change during the bath-application of ACh (30 µM) recorded from the same neuron pairs shown in (A,B).

### Supplemental Tables

### Table S1

| | Statistical significance | | | |
|---|---|---|---|---|
| | L6a FOXP2+ vs L6a FOXP2− | L6b FOXP2+ vs L6b FOXP2− | L6a FOXP2+ vs L6b FOXP2+ | L6a FOXP2− vs L6b FOXP2− |
| ***Soma*** | | | | |
| Depth (μm) | 0.089 | 0.74 | **$2.88\times10^{-3}$** | **$2.06\times10^{-4}$** |
| Relative depth (%) | 0.083 | 0.14 | **$1.08\times10^{-5}$** | **$1.08\times10^{-5}$** |
| ***Dendrite*** | | | | |
| No. of apical dendrite branches | **$1.94\times10^{-3}$** | N/A | 0.30 | N/A |
| Total length of apical dendrite (μm) | **$8.93\times10^{-3}$** | N/A | 0.39 | N/A |
| No. of basal dendrites | 0.36 | 0.098 | 0.42 | 1.06 |
| No. of basal dendrite branches | 0.18 | **$6.30\times10^{-3}$** | 0.93 | **0.03** |
| Total length of basal dendrites (μm) | **$2.88\times10^{-3}$** | **$2.17\times10^{-5}$** | 0.58 | **$2.88\times10^{-3}$** |
| Horizontal field span of dendrite (μm) | **0.019** | **$1.08\times10^{-5}$** | 0.35 | **$2.06\times10^{-4}$** |
| No. of dendritic branches | **0.045** | 0.14 | 0.42 | 0.34 |
| Total length of dendrites (μm) | 0.68 | 0.58 | 0.31 | 0.91 |
| ***Axon*** | | | | |
| Horizontal field span of axon (μm) | **$1.08\times10^{-5}$** | **$4.33\times10^{-5}$** | **0.019** | 0.58 |
| No. of axonal branches | **$7.80\times10^{-4}$** | **$1.40\times10^{-3}$** | 0.45 | **0.037** |
| Total length of axon (μm) | **$4.33\times10^{-5}$** | **$3.25\times10^{-4}$** | 0.44 | **0.011** |

**Table S1.** Statistical tests for the morphological properties of L6a and L6b FOXP2+ and FOXP2− excitatory neurons. P value was calculated using the non-parametric Wilcoxon-Mann-Whitney two-sample rank test between individual groups.

### Table S2

| | Statistical significance | | | |
|---|---|---|---|---|
| | L6a FOXP2+ vs L6a FOXP2− | L6b FOXP2+ vs L6b FOXP2− | L6a FOXP2+ vs L6b FOXP2+ | L6a FOXP2− vs L6b FOXP2− |
| ***Passive*** | | | | |
| Vrest (mV) | 0.91 | 0.39 | **0.02** | **$1.59\times10^{-3}$** |
| Rin (MΩ) | 0.058 | 0.75 | **$5.54\times10^{-3}$** | **$9.51\times10^{-4}$** |
| Tau (ms) | **0.021** | 0.14 | **$1.47\times10^{-3}$** | 0.16 |
| Sag (%) | 0.24 | **$4.91\times10^{-5}$** | 0.34 | **$9.72\times10^{-3}$** |
| ***Single AP*** | | | | |
| Rheobase current (pA) | 0.89 | 0.32 | **$2.71\times10^{-3}$** | **$1.22\times10^{-3}$** |
| AP threshold (mV) | 0.16 | 0.98 | 0.11 | 0.91 |
| AP half-width (ms) | **$2.19\times10^{-8}$** | 0.32 | **$8.11\times10^{-11}$** | **$9.72\times10^{-3}$** |
| AP amplitude (mV) | 0.96 | 0.58 | **0.017** | 0.40 |
| AP latency (ms) | **$1.05\times10^{-6}$** | **$2.23\times10^{-3}$** | **$6.21\times10^{-4}$** | 0.24 |
| fAHP amplitude (mV) | **$1.47\times10^{-4}$** | 0.08 | 0.07 | 0.077 |
| fAHP latency (ms) | **$6.60\times10^{-5}$** | 0.94 | **$4.51\times10^{-11}$** | **$7.59\times10^{-3}$** |
| ***Repetitive firing*** | | | | |
| Max. firing frequency (Hz) | 0.24 | 0.69 | **0.042** | 0.32 |
| Slope of F-I curve (APs/100 [illegible] | 0.70 | 0.88 | **0.046** | 0.16 |
| Adaptation ratio | 0.31 | 0.22 | 0.66 | 0.70 |
| s.d. of ISIs (ms) | **$7.45\times10^{-5}$** | 0.32 | **$2.64\times10^{-4}$** | 0.057 |
| ISI1 (ms) | 0.12 | **0.025** | **$2.48\times10^{-4}$** | **$2.67\times10^{-6}$** |
| ISI2 (ms) | **$5.0\times10^{-3}$** | **0.024** | 0.23 | 0.092 |
| ISI3 (ms) | 0.78 | **0.034** | 0.52 | 0.13 |
| AP1 amplitude (mV) | 0.52 | 0.37 | 0.10 | 0.46 |
| AP2 amplitude (mV) | **$4.87\times10^{-7}$** | 0.11 | **$1.6\times10^{-3}$** | **$6.89\times10^{-4}$** |
| AP3 amplitude (mV) | **0.028** | 0.43 | **0.021** | **0.013** |
| AP1 half-width (ms) | **$1.06\times10^{-8}$** | 0.35 | **$1.13\times10^{-10}$** | **0.02** |
| AP2 half-width (ms) | **$7.64\times10^{-7}$** | **$2.16\times10^{-3}$** | **$1.14\times10^{-10}$** | 0.36 |
| AP3 half-width (ms) | **$2.09\times10^{-10}$** | **$8.07\times10^{-3}$** | **$1.13\times10^{-10}$** | 0.60 |
| AP1 threshold (mV) | 0.099 | 0.51 | 0.39 | 0.97 |
| fAHP1 amplitude (mV) | **$5.79\times10^{-8}$** | 0.15 | 0.31 | **$2.04\times10^{-3}$** |
| AP9 threshold (mV) | 0.32 | 0.75 | 0.31 | 0.84 |
| fAHP9 amplitude (mV) | 0.36 | 0.33 | **$7.60\times10^{-3}$** | **$9.09\times10^{-3}$** |
| fAHP9 - fAHP1 (mV) | **$6.53\times10^{-11}$** | 0.055 | **0.035** | **$7.10\times10^{-5}$** |

**Table S2.** Statistical tests for the electrophysiological properties of L6a and L6b FOXP2+ and FOXP2− excitatory neurons. P value was calculated using the non-parametric Wilcoxon-Mann-Whitney two-sample rank test between individual groups.

**Table S3**

| | **L6b FOXP2– inverted (n=17)** | **L6b FOXP2– multipolar (n=13)** | **p value** |
|---|---|---|---|
| ***Passive*** | | | |
| $V_{rest}$ (mV) | -68.6 ± 6.8 | -71.5 ± 6.0 | 0.28 |
| $R_{in}$ (MΩ) | 221.5 ± 59.5 | 187.9 ± 37.1 | **0.039** |
| $\tau_M$ (ms) | 26.5 ± 5.6 | 21.2 ± 6.3 | **0.017** |
| Sag (%) | 7.9 ± 2.7 | 4.5 ± 3.9 | **0.0091** |
| ***Single AP*** | | | |
| Rheobase current (pA) | 96.5 ± 41.4 | 99.2 ± 33.2 | 0.61 |
| AP threshold (mV) | -36.0 ± 4.1 | -40.7 ± 3.9 | **0.008** |
| AP half-width (ms) | 1.02 ± 0.16 | 1.15 ± 0.22 | **0.043** |
| AP amplitude (mV) | 89.0 ± 6.8 | 97.7 ± 8.5 | **0.0045** |
| AP latency (ms) | 117.6 ± 49.1 | 302.8 ± 125.8 | **$3.41 \times 10^{-5}$** |
| fAHP amplitude (mV) | 5.5 ± 3.5 | 11.9 ± 2.9 | **$2.00 \times 10^{-5}$** |
| fAHP latency (ms) | 4.3 ± 0.9 | 24.5 ± 15.3 | **$5.00 \times 10^{-7}$** |
| ***Repetitive firing*** | | | |
| Max. firing frequency (Hz) | 26.0 ± 8.2 | 19.5 ± 5.4 | **0.012** |
| Slope of F-I curve (APs/ 100 pA) | 18.1 ± 4.5 | 9.2 ± 4.4 | **$7.54 \times 10^{-5}$** |
| Adaptation ratio | 1.1 ± 0.1 | 2.1 ± 1.0 | **$2.50 \times 10^{-5}$** |
| s.d. of ISIs (ms) | 7.3 ± 3.0 | 36.4 ± 17.9 | **$3.85 \times 10^{-8}$** |
| ISI1 (ms) | 11.7 ± 9.7 | 32.6 ± 10.7 | **$3.41 \times 10^{-5}$** |
| ISI2 (ms) | 56.2 ± 33.0 | 43.9 ± 11.5 | 0.30 |
| ISI3 (ms) | 101.2 ± 22.1 | 52.1 ± 11.3 | **$1.66 \times 10^{-8}$** |
| AP1 amplitude (mV) | 88.3 ± 6.4 | 96.1 ± 7.5 | **0.0039** |
| AP2 amplitude (mV) | 64.0 ± 9.5 | 91.2 ± 8.8 | **$1.17 \times 10^{-7}$** |
| AP3 amplitude (mV) | 77.4 ± 7.8 | 90.2 ± 8.7 | **$4.95 \times 10^{-4}$** |
| AP1 half-width (ms) | 0.96 ± 0.14 | 1.06 ± 0.23 | 0.094 |
| AP2 half-width (ms) | 1.64 ± 0.36 | 1.41 ± 0.38 | 0.10 |
| AP3 half-width (ms) | 1.47 ± 0.32 | 1.44 ± 0.40 | 0.80 |
| AP1 threshold (mV) | -36.9 ± 4.0 | -41.0 ± 5.0 | **0.016** |
| fAHP1 amplitude (mV) | 2.3 ± 3.9 | 6.3 ± 2.5 | **$7.34 \times 10^{-4}$** |
| AP9 threshold (mV) | -32.5 ± 4.5 | -38.1 ± 5.4 | **0.0013** |
| fAHP9 amplitude (mV) | 12.5 ± 2.1 | 10.3 ± 1.7 | **0.0051** |
| fAHP9 - fAHP1 (mV) | -5.84 ± 2.72 | -1.11 ± 2.30 | **$1.62 \times 10^{-4}$** |

**Table S3**. Statistical comparison of the electrophysiological properties of inverted and multipolar L6b FOXP2– excitatory neuron subtypes. p value was calculated using the non-parametric Wilcoxon-Mann-Whitney two-sample rank test.
